# Sex differences in thyroid aging and their implications in thyroid disorders: insights from gene regulatory networks

**DOI:** 10.1101/2025.10.27.684890

**Authors:** Enakshi Saha

**Affiliations:** Department of Epidemiology and Biostatistics, University of South Carolina Arnold School of Public Health, Columbia, SC 29208, USA

## Abstract

Thyroid disorders are more prevalent in females than males, with sex differences in disease risk varying by age, e.g. although risk of anaplastic thyroid carcinoma (ATC) increases with age for both sexes, females are diagnosed at younger ages than males. In contrast, Hashimoto’s thyroiditis (HT) is more common in ages 30-50, with females having higher risk. These patterns suggest that in thyroid, gene regulatory processes evolve with age in a sex-specific manner. To characterize these sex differences in thyroid aging, we estimated and analyzed individual-specific gene regulatory networks using BONOBO and PANDA algorithms. We found that regulatory patterns of pathways involved in cell proliferation, immune response, and metabolic processes vary nonlinearly with age, with sex-specific differences in direction and rate. Females show two inflection points around ages 40 and 60, when regulatory patterns show significant change for most pathways, consistent with changes in targeting by estrogen and androgen receptors. Investigating gene regulatory patterns in HT and ATC, we observed that disease-relevant immune and metabolic pathways changed with age in the same direction as they change in ATC. In contrast, in HT, disease-related regulatory patterns are in the opposite direction of aging. In age groups where HT is most diagnosed, regulatory patterns in females more closely resembled disease states than males, highlighting sex-biased regulation’s role in elevated disease risk in females. Our findings demonstrate immune response and metabolic processes are regulated in age- and sex-specific manner, potentially driving variation in disease risk, emphasizing the need for tailored screening and prevention strategies.

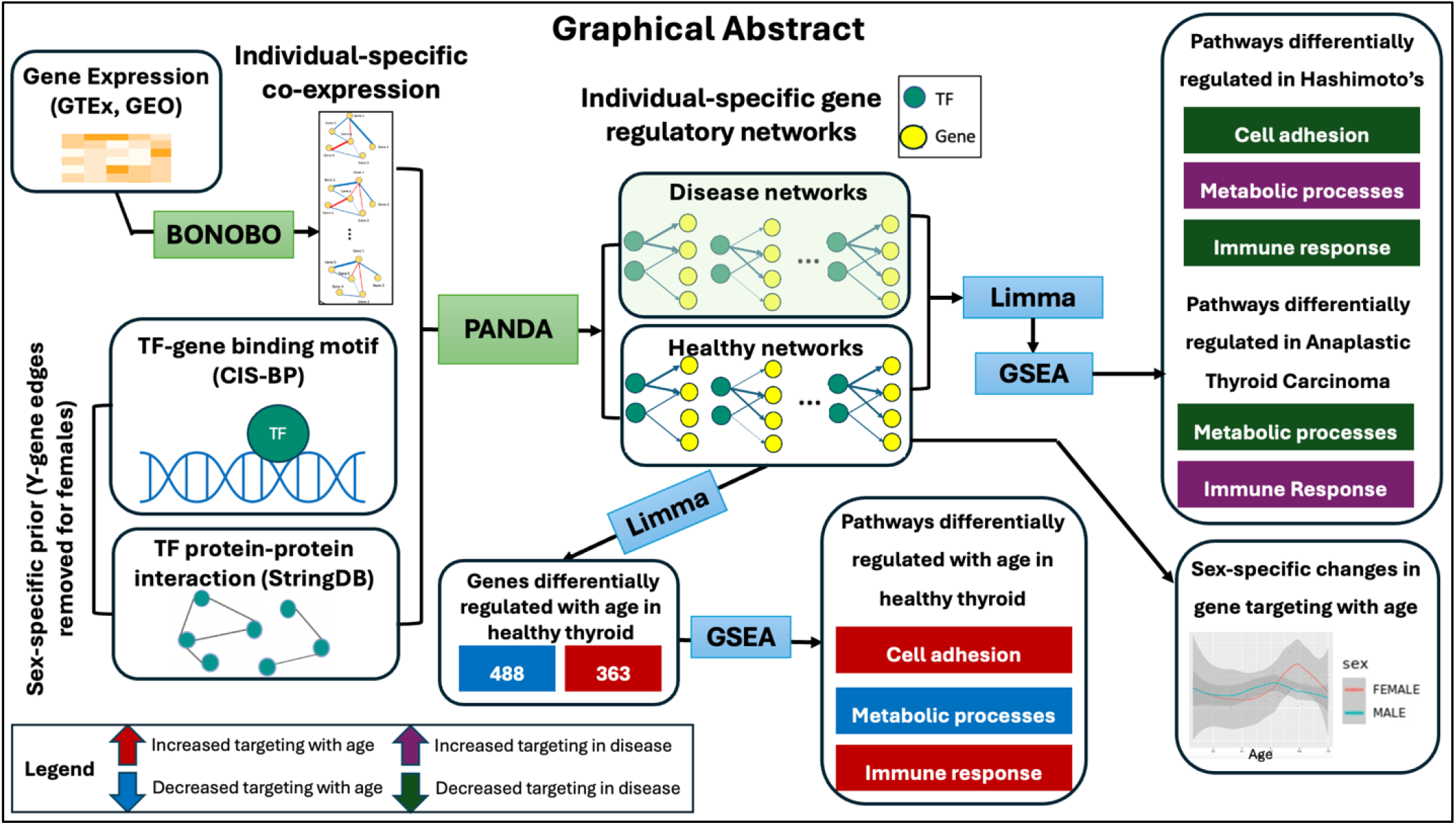

## Introduction

Thyroid disorders, including autoimmune diseases such as Hashimoto’s thyroiditis (HT) and thyroid cancer subtypes such as Anaplastic Thyroid Carcinoma (ATC) are more prevalent in females than in males^1^. However, this sex difference in disease risk is not uniform across all age groups. For example, while the risk of thyroid cancers increases with age in males, many thyroid cancer subtypes including papillary and follicular thyroid carcinomas disproportionately affect females below age 50, compared to females of older age groups^2^. Thyroid conditions such as hypothyroidism, autoimmune diseases and thyroid cancers not only affect quality of life of the afflicted individuals, but also exacerbate pre-existing conditions such as cardiovascular disease^3^, depression^4^, and infertility^5^; thyroid conditions also increase the risk of several other diseases including metabolic syndrome^6^, osteoporosis^7^, cognitive impairment^8^, and cancers of breast, lung and prostate^9^. Understanding the molecular mechanisms driving age- and sex-specific disparities in thyroid disorders is important to mitigate disease risk across diverse subpopulations.

The influence of sex steroid hormones has been cited as a potential driver of higher thyroid disease risk in females^10,11^, which in turn suggests that variations in sex steroid hormone levels in males and females over lifetime might be a contributing factor behind an age-dependent sex-biased disease rate. Among other factors driving age- and sex-biased thyroid disease risk, studies have found sex-specific regulation of immune cells^12^, skewed X chromosome inactivation leading to a sex-biased dosage of X-linked gene expression^13^, and sex-differences in DNA methylation patterns^14^. However, these factors alone offer only a fragmented understanding of disease etiology, and there is a need for a systems-based approach looking into the interplay between genes, their molecular regulators such as transcription factors (TFs) and their combined influence on the immune system and sex-hormone receptors. Such an approach will facilitate a deeper understanding of the biological mechanisms driving the risk of thyroid disorders, how these are affected by sex, and how they change over the life course. Gene regulatory networks (GRN) are one such systems-based approach; GRNs are a powerful tool for exploring how the interactions between genes and regulatory TFs vary in ways that alter biological processes to drive complex diseases^15–19^ including cancer.

We estimated individual sample-specific gene regulatory networks for normal thyroid tissue, as well as for individuals with Hashimoto’s thyroiditis (HT) and anaplastic thyroid carcinoma (ATC) using various public databases including the Genotype Tissue Expression Project^20^ (GTEx) and the Gene Expression Omnibus^21^ (GEO). HT is an autoimmune condition that is most frequently diagnosed in individuals between ages 30 and 50; ATC is an aggressive form of thyroid cancer that is more commonly diagnosed in adults above the age of 50. Both diseases are known to disproportionately affect females.

To estimate these networks, we used BONOBO^22^ and PANDA^23^ which together can be used to infer individual-specific gene regulatory networks, consisting of weighted edges between genes and the TFs that regulate them. BONOBO is an empirical Bayesian model that infers individual-specific gene-gene co-expression networks. These individual-specific co-expression networks are used as inputs to a message passing algorithm PANDA, which integrates them with prior knowledge on TF-gene motif information and TF-TF interactions (from protein-protein interaction data), to estimate individual-specific TF-gene regulatory networks. The resulting networks are represented as bipartite graphs of regulatory relationships between TFs and genes, where the network edge weights represent the strength of the estimated regulatory associations between genes and TFs. These Individual-specific networks capture regulatory interactions within each subject, enabling us to analyze how gene regulatory relationships vary by age and sex across the population.

To quantify regulatory activities in these networks, we defined a “targeting score” as the sum of the weights of all edges associated with each TF (out-degree) or gene (in-degree). In normal thyroid tissue, we identified TFs and genes whose targeting scores varied by sex and age. We then compared these patterns with those in HT and ATC networks to understand how age- and sex-biased regulatory differences relate to disease risk. Differentially targeted genes were analyzed for biological process enrichment, and key findings were validated using molecular data from GEO and hormonal data from the National Health and Nutrition Examination Survey (NHANES).^24^

Our findings indicate a potential role for TFs, including estrogen and androgen receptor TFs, in driving HT at younger ages and ATC at older ages as well as contributing to the greater risk observed in females compared to males. Further, we find that aging in thyroid is a sex-biased process and that the associated age- and sex-biased gene regulatory differences may contribute to differential risk of thyroid conditions.

## Results

### Gene Regulatory Networks Vary with Age in a Sex-biased Manner

We constructed individual-specific gene regulatory networks for 653 normal thyroid samples from GTEx (434 males, 219 females) using a two-step approach (**Figure 1 A**). First, we generated individual-specific gene-gene co-expression networks using BONOBO. Second, we integrated these with sex-specific TF-gene motif priors and TF protein-protein interaction (PPI) data using the PANDA message-passing algorithm. For each network, we computed gene targeting scores (in-degrees), defined as the sum of all edge weights from regulatory TFs to each gene, quantifying the total regulatory input that gene receives (**Figure 1 B**). We hypothesized that genes with targeting scores differentially changing by age and sex would be associated with disease susceptibility. Detailed methods for constructing sex-specific priors and individual-specific networks are provided in *Methods*.

**Figure 1:**
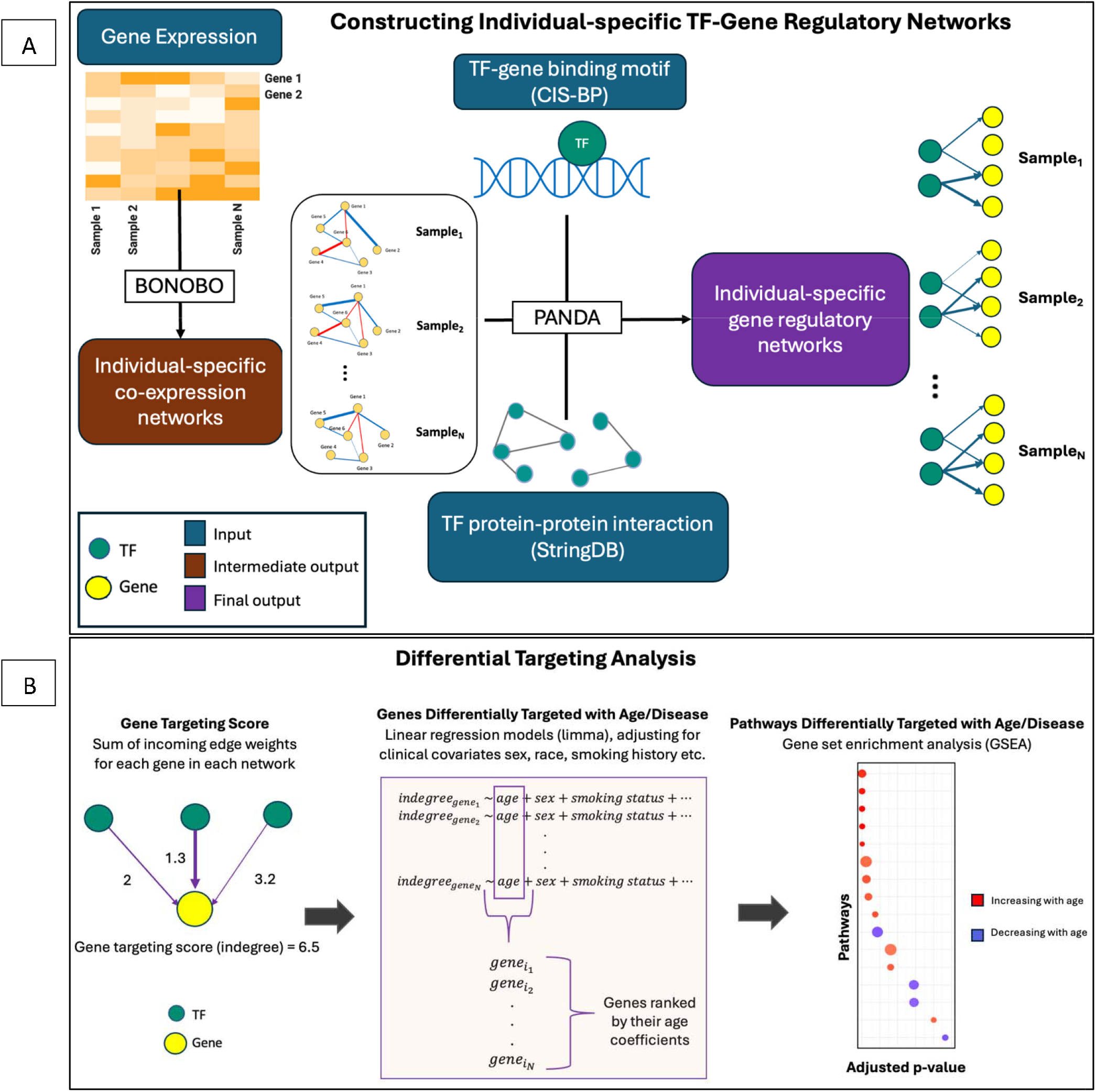
Overview of the study. (A) Constructing individual-specific TF-gene regulatory networks using BONOBO+PANDA. Individual-specific co-expression networks computed using BONOBO are used as inputs to PANDA, along with TF-gene motif and PPI networks to estimate individual-specific TF-gene regulatory networks. (B) Overview of the differential targeting analysis.

We used linear models for gene-in-degrees on age and adjusted for clinical covariates (sex, race, BMI, smoking status, ischemic time, RNA integrity number and batch), which allowed us to identify (**Figure 2 A**) a total of 751 genes (henceforth referred to as the “aging genes”, Supplementary Table S0) that are significantly (p-value < 0.05) differentially targeted by TFs with age. Among these genes, 363 genes were observed to be increasingly targeted with age, and the remaining 388 genes were decreasingly targeted with age by TFs.

**Figure 2:**
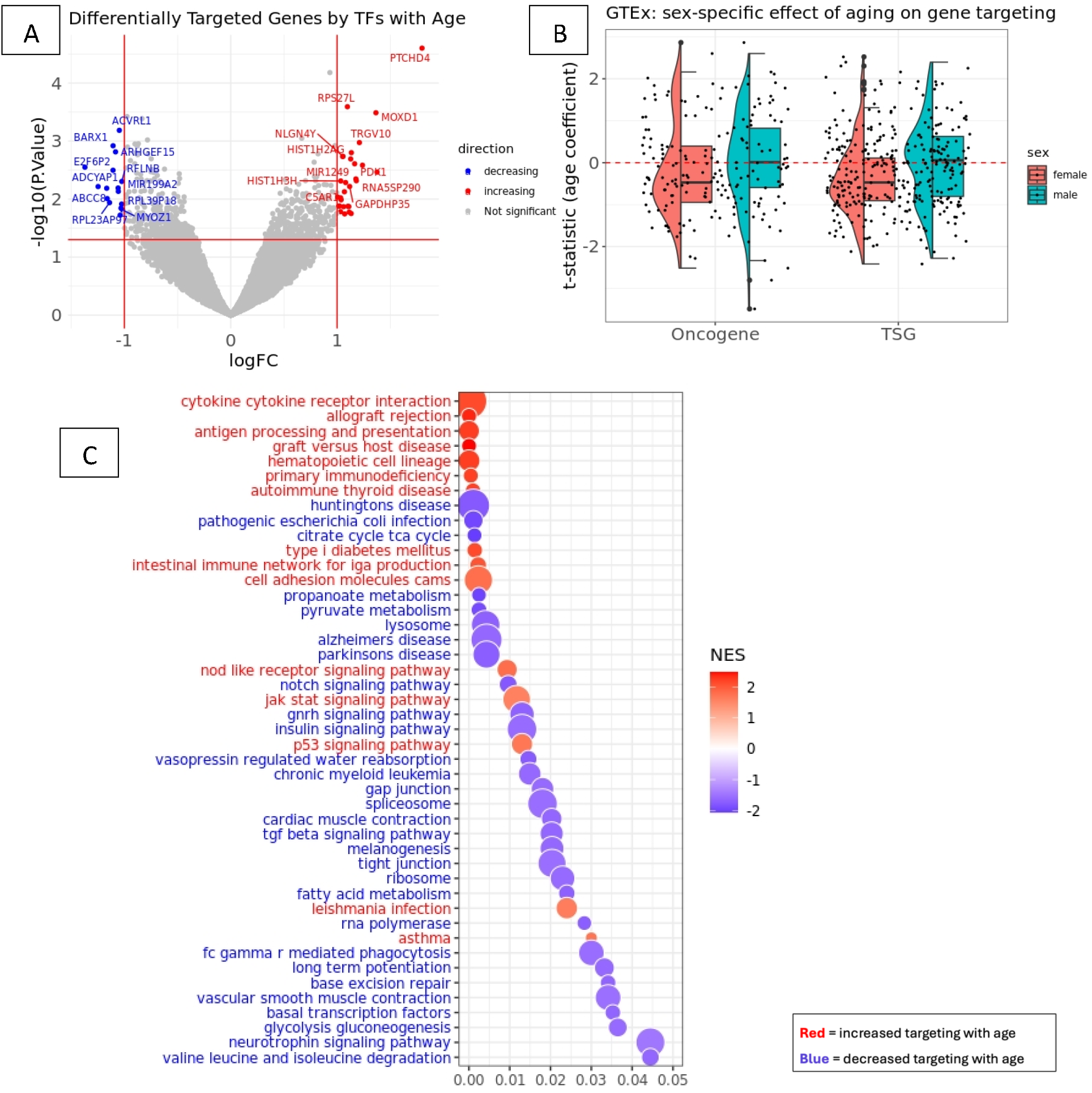
Gene regulatory changes associated with age. (A) Volcano plot of genes that are differentially (increasingly or decreasingly) targeted by transcription factors (TFs) over varying age in normal thyroid tissue samples from GTEx. The x-axis represents log fold change (logFC), defined as the change in gene in-degree with a unit change in age. The y-axis represents negative logarithm of p-values (-log10(P.Value)). (B) Boxplot of the t-statistics of the age coefficients from the limma models in GTEx, corresponding to the oncogenes and tumor suppressor genes (TSG) from the COSMIC database. Genes with positive (and negative) values of t-statistics are targeted more (and less) by TFs with age. (C) Bubble plot showing the normalized enrichment scores (NES) of biological pathways that are significantly (false discovery rate-adjusted p-value < 0.05) increasingly (marked red) or decreasingly (marked blue) targeted by TFs with age. NES are obtained from ranked gene set enrichment analysis. The x-axis shows the adjusted p-values, and the y axis shows the names of the pathways. Bubble size is proportional to the number of genes in specific pathways.

Many of these genes are known to play important roles both in aging and various thyroid diseases. These include *ACE2*^*25*^, *TNFAIP3*^*26*^, *PTPN22*^*27*^, and *HLA-DQA1*^*28*^, which are implicated in autoimmune thyroid diseases including HT and Grave’s disease. Some genes are implicated in thyroid cancer, with *TNFAIP3*^*29*^ also exhibiting tumor suppressive functions. Additionally, genes relevant to thyroid cancer including proto-oncogene *CXCR4*^*30*^, and TSG *PTPRC*^*31*^ and *SMAD3*^*32*^ were differentially targeted with age in normal thyroid. However, it is worth noting that although the TF-targeting patterns of these genes vary with age in both males and females, there is considerable difference between sexes in how TF-targeting of specific genes varies with age in general.

To systematically examine these sex-specific age-related patterns in cancer-relevant genes, we performed a comprehensive analysis (**Figure 2 B**) of TF-targeting patterns of proto-oncogenes and tumor suppressor genes (TSG) catalogued in the somatic mutations in cancer (COSMIC) database^33^. For the 94 oncogenes and 171 TSGs recorded in the database, we found that, on average, oncogenes exhibited significantly decrease in targeting with age in females, as evidenced by a one-sided Wilcoxon signed-rank test on the t-statistics for age coefficients of these genes (p-value =0.0297). In contrast, in males, the targeting pattern of oncogenes did not change significantly with age in any direction (p-value =0.8771 for two-sided test). We found similar patterns for TSGs as well (p-value = 2.818e-08 and 0.8744, for a one-sided test for females and a two-sided test for males, respectively), indicating an age-related decline in tumor suppressive gene regulatory mechanisms in females, but not in males. For reference, for the other genes, those not in the COSMIC database, there was an overall decline in gene targeting with age in females (p < 2.2e-16 for one-sided test) while for males we found an overall increase in gene targeting with age (p-value = 6.86e-09 for one-sided test). Within females, although the age-related decline in oncogene targeting was in line with non-cancer-related genes, targeting of TSGs declined at a faster rate (p-value = 0.01582), as evidenced by a two-sample two-sided Wilcoxon signed rank test between the t-statistics of the age-coefficients for TSGs and non-cancer-related genes. Test directionality (one-sided vs. two-sided) was determined based on visual inspection of the data (**Figure 2 B**).

For networks derived from the GTEx data, we performed a ranked gene set enrichment analysis (Supplementary Table S1) based on the age coefficients from the linear model on gene in-degrees and identified several biological pathways that were significantly (p-value < 0.05 after FDR correction) differentially targeted by TFs as a function of age (**Figure 2 C**). These pathways were validated in an independent dataset of healthy thyroid samples (GEO accession number GSE165724; Supplementary Table S2) (**Figure S1**) using the same differential targeting analysis procedure as in GTEx. We found that several key pathways involved in immune response and immune-related diseases such as cytokine-cytokine receptor interaction, hematopoietic cell lineage, intestinal immune network for Iga production, and the pathway associated with autoimmune thyroid disease were significantly increasingly targeted by TFs with age. A cell type deconvolution analysis (**Figure S2**) using “xcell”^34^ showed that this difference in the regulation of immune pathways was not correlated with variations in immune cell proportion in thyroid tissue, as the proportion of immune cells did not change with age in females, even though among males, the proportion of CD4+ Th2 helper T cells increased with age while the proportion of T cell gamma delta and common myeloid progenitors decreased with age.

Pathways involved in metabolic processes including propanoate metabolism, fatty acid metabolism, and insulin signaling pathway were significantly (p-value < 0.05 after FDR correction) decreasingly targeted by TFs with age. Targeting of pathways associated with cell adhesion, cell migration and proliferation, including the tight junction pathway and the focal adhesion pathway, also declined with age, while targeting of the cell adhesion molecules (CAMs) increased with age. The notch signaling pathway, a cell signaling pathway associated with thyroid cancer^35^, was decreasingly targeted by TFs with age. In contrast, the P53 signaling pathway, which has been shown to hinder progression of thyroid cancers to more aggressive states^36^, showed increased targeting by TFs with age.

Upon closer inspection of TF-targeting of all genes in specific groups of pathways across individuals of different ages, we found that the aging-related alterations in the TF-targeting of biological processes and pathways are non-linear, meaning that gene targeting by TFs does not increase or decrease at a uniform rate with age (**Figure 3**). Moreover, the targeting patterns differed between males and females over the entire age range considered. For example, within females, we found that for most pathways, including those involved in immune system, cell signaling, cell growth and death, and the endocrine system, there were two inflection points around the ages 40 and 60, where TF targeting patterns changed rapidly. Given that the average age of menopause reported in the USA is 52^37^, these inflection points in this age group in TF-targeting of key biological pathways indicate substantial physiological changes at the molecular level, accompany other menopausal changes. In males, we observed milder inflection points around the same age range that affect pathways associated with immune response, immune diseases, cell proliferation and cell signaling. However, the male-specific inflection points were less pronounced compared to the gene regulatory shifts we observed in females of similar age.

**Figure 3:**
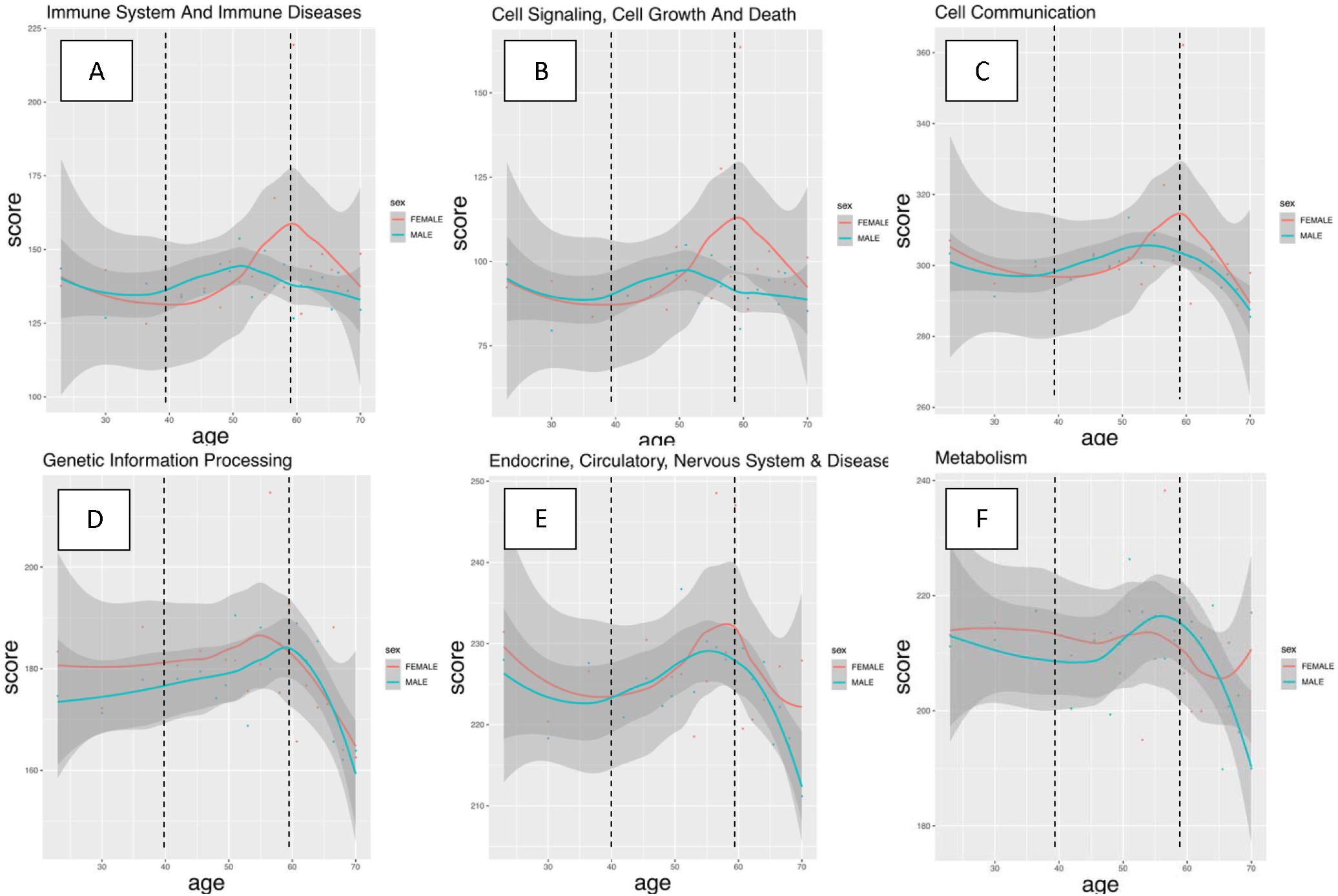
Sex-specific variations in pathway targeting scores with age in normal thyroid. The six plots (A-D) show how the targeting scores change with age in each sex for six groups of biological pathways (grouped according to KEGG hierarchy of pathways). The pathway targeting score for a specific group of pathways is the sum of in-degrees of all genes in that group of pathways. The x-axis represents age, and the y-axis represents the gene targeting score of the corresponding group of pathways. Each plot shows an aging trajectory curve computed for male samples (marked teal) and a trajectory curve computed for female samples (marked reddish orange) in GTEx using Locally Estimated Scatterplot Smoothing (LOESS). The grey shaded region around each curve is the pointwise 95% confidence band.

We performed sensitivity analyses using three complementary approaches and concluded that the observed non-linear TF-targeting trajectories of biological processes in GTEx described above, were robust to the choice of smoothing parameter (**Figure S4**), sample composition across age groups (**Figure S5**), and the choice of smoothing method (**Figure S6**).

### Sex Hormone Receptor Targeting Associated with Marked Shift in Regulatory Patterns Around Middle Ages in Females

Androgen (AR) and estrogen receptors (ESR1 and ESR2) play important role in aging^38^ and disease^39^ by acting as TFs that mediate hormone-dependent gene regulation. In females, we found that the age-associated regulatory edges connecting the 751 aging genes with sex hormone receptor TFs, including ESR1, ESR2 and AR weaken and sharply decline between ages 40 and 60 (**Figure 4 A-C**), thus suggesting that the shift in regulatory targeting of key biological pathways between ages 40 and 60 in females is associated with declining activity of sex hormone receptor TFs. Our analysis on the sex hormonal levels on the independent dataset NHANES^24^ demonstrated that this pattern is consistent with a decline in estradiol levels in females around the same age (**Figure S3**). In contrast, within males, the sex hormone receptor TFs did not show any considerable age-dependence in gene targeting (**Figure 4 A-C**). This is supported by the relative stability of the sex hormonal levels with age in males, reported in the NHANES data (**Figure S3**).

**Figure 4:**
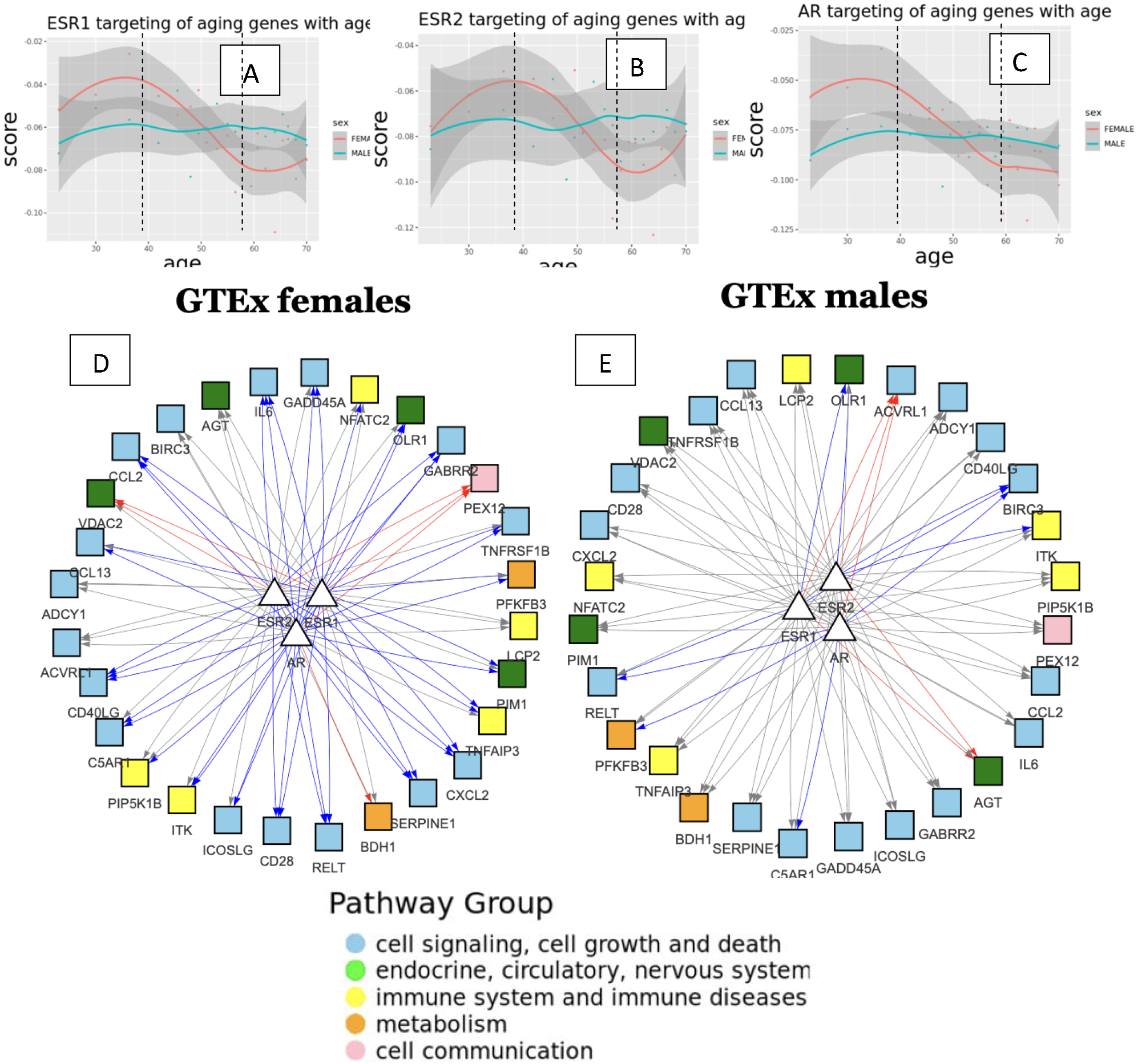
Sex-specific change in transcription factor (TF) out-degree with age in normal thyroid. (A-C) The plot shows TF out-degree (sum of all outgoing edge weights from the TF to the aging-related genes) of (A) estrogen receptor 1 (ESR1), (B) estrogen receptor 2 (ESR2) and (C) androgen receptor (AR). The x-axis shows age, and the y-axis shows targeting score or the TF out-degree. The grey shaded region around each curve is the pointwise 95% confidence band. (D-E) Subnetwork of the most significant aging-associated edges connected to ESR1, ESR2 and AR in (D) females and (E) males. Edges marked red and blue significantly (p-value < 0.05) increase and decrease in weight with age, respectively. Edges that do not change significantly with age are marked grey. TFs are represented by white triangles and genes are represented by colored squares. Gene nodes are marked by their pathway membership in the KEGG database.

Upon closer inspection of the neighborhood of ESR1, ESR2 and AR in the gene regulatory networks in females (**Figure 4 D**), this aging-related changes in gene targeting by sex hormone receptors is most pronounced for genes associated with cell signaling, cell growth and death, and immune response. However, as can be seen in males (**Figure 4 E**), most edges connecting ESR and AR TFs with their target genes do not change significantly (p-value > 0.05) with age.

### Aging-associated Pathways are Differentially Targeted by TFs in Thyroid Diseases

To investigate how aging-related regulatory changes contribute to disease susceptibility, we analyzed the implications of age- and sex-biased gene regulatory patterns in thyroid in the context of two thyroid disorders, both of which are more common in females than males: Hashimoto’s thyroiditis (HT), an autoimmune condition more prevalent in younger adults below age 50 and Anaplastic Thyroid Carcinoma (ATC), an aggressive form of thyroid cancer, which primarily affects older adults above age 50.

For each of the two diseases, we compared individual-specific gene regulatory networks derived from normal thyroid tissue with those derived from the corresponding diseased tissue samples and identified several genes and biological pathways that were differentially targeted by TFs in diseases. In analyzing individuals in GTEx with HT, we identified 32 genes (FDR-corrected p-value < 0.05) that were more highly targeted in individuals with HT than those without, including *CALR*, an immune-related gene over-expressed in HT^40^; these were validated (same effect direction with FDR-corrected p-value < 0.05) in an independent dataset, GEO accession number GSE29315. We also identified 197 genes (discovered in GTEx and validated in GSE29315) that were targeted at lower levels in individuals with HT, compared to healthy individuals, including immune-related genes *CTLA-4* and *PTPN22*, both known drivers of HT risk^41^.

In analyzing ATC gene regulatory networks from a dataset with GEO accession number GSE33630, we found 301 genes that were significantly (FDR corrected p-value < 0.05) more highly targeted in tumor than in normal tissue, including *AKAP4*, the silencing of which has been shown to inhibit thyroid cancer progression^42^ and *CEBPB*, the over-expression of which is reported to drive poorer prognosis in ATC^43^. Among other ATC-related genes, we found 670 genes to have significantly (FDR-corrected p-value < 0.05) reduced targeting in tumor, compared to normal tissue, including immune-related genes *FOS* and *JUN*, both of which have been reported as potential diagnostic markers in thyroid cancer^44^. These genes were validated (same effect direction with FDR-corrected p-value < 0.05) in an independent dataset with GEO accession number GSE27155.

For each disease, we performed gene set enrichment analysis to identify biological processes and pathways that were differentially targeted by TFs in disease. For both diseases, we were able to validate (same effect direction with FDR-corrected p-value < 0.05) most of the disease-associated pathways in independent datasets (with GEO accession numbers GSE29315 for HT and GSE27155 for ATC). It is noteworthy that many of these disease-specific pathways were among those that we had found to be differentially targeted with age (**Figure 5**) in normal thyroid tissue. For example, we found that TF-associated regulatory targeting of metabolic pathways such as fatty acid metabolism, and pyruvate metabolism were significantly higher (FDR-corrected p-value < 0.05) within individuals with HT, than in those without. These same metabolic pathways were decreasingly targeted by TFs with age in regulatory networks from normal thyroid samples in GTEx. In contrast, pathways associated with immune response such as cytokine-cytokine receptor interaction, hematopoietic cell lineage, intestinal immune network for Iga production, and also the pathway associated with autoimmune thyroid disease, were increasingly targeted with age in normal thyroid, and exhibited lower targeting in HT, compared to normal tissue, meaning that aging-associated gene regulatory patterns in normal tissue are in the opposite direction to the disease-related gene regulatory patterns in HT. Thus, regulatory interaction patterns between TFs and their target genes that resemble those in younger individuals (below age 50) may contribute to the higher incidence of the disease among individuals aged 30–50 compared to older adults.

**Figure 5:**
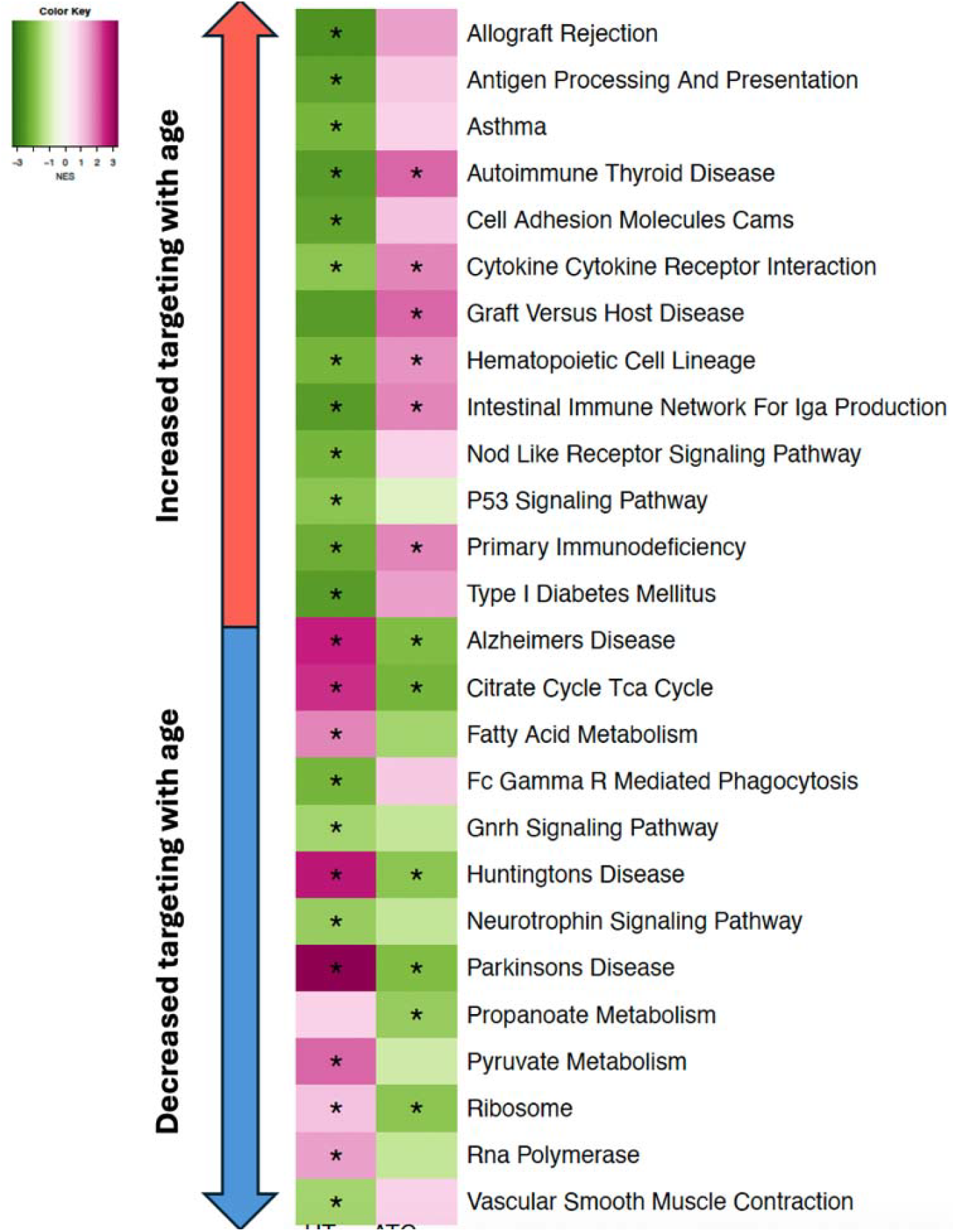
Pathways that are significantly differentially targeted by TFs with age and also in Hashimoto’s thyroiditis (HT) and/or Anaplastic Thyroid Carcinoma (ATC). The heatmap represents pathways in rows and disease in columns. Intensity of (green and pink) colors are proportional to the normalized enrichment scores (NES) from the gene set enrichment analysis (GSEA) performed to identify disease-related pathways, separately for HT and ATC compared to GTEx healthy thyroid. Pathways targeted more in disease compared to normal tissue are colored pink and pathways targeted less in disease compared to normal tissue are colored green. Pathways that were significant (p-value 0.05, after adjusting for false discovery rate) are marked with an asterisk. The left bar shows red or blue beside the pathway if that pathway is increasingly or decreasingly targeted by TFs with age, respectively.

In ATC, pathways involved in immune response, including autoimmune thyroid disease, cytokine-cytokine receptor interaction, intestinal immune response for IgA production and primary immunodeficiency, were more highly targeted by TFs, compared to the targeting in normal thyroid.

Metabolic pathway genes, including those in citrate cycle TCA cycle and propanoate metabolism pathways had lower regulatory targeting in ATC than normal thyroid. Earlier we observed similar TF-targeting patterns for aging, where immune pathways were increasingly targeted by TFs as a function of age and metabolic pathways were decreasingly targeted with age.

Overall, we found that in ATC, disease-associated TF targeting patterns across most pathways resembled those observed with aging, whereas in HT, TF targeting showed the opposite trend relative to age-associated changes. This supports the hypothesis that individual-specific regulatory patterns may be a contributing factor to the higher incidence of ATC in older adults and the higher incidence of HT diagnosis in those below age 50.

The complete GSEA results from GTEx samples with HT, GSE29315 samples with HT, GSE33630 samples with ATC and GSE27155 samples with ATC are provided in the Supplementary Table S3-S6 respectively.

### Females Show Enhanced Disease-Associated Gene Regulatory Signatures in High-Risk Age Groups

We also explored whether the TF-targeting patterns of disease-associated genes change with age in a sex-biased manner, to understand why females have disproportionately higher risk of most thyroid diseases, than do males. For each disease, we identified genes that had significantly (p-value <0.05) either higher or lower TF-targeting in disease, compared to normal tissue, in the discovery datasets (GTEx samples with HT and dataset with GEO accession number GSE33630 for ATC). From these genes, we selected those that were validated in the regulatory networks derived from the respective independent validation datasets (GEO accession numbers GSE29315 for HT and GSE27155 for ATC), and divided them into two gene sets for each disease (four gene sets in total) depending on whether they were either more or less targeted by TFs in the disease states relative to normal thyroid.

For each gene set, we defined a gene-set score for each individual as the sum of incoming edge weights from TFs to all genes on the set based on the corresponding individual-specific gene regulatory network. We also computed the four gene set scores (two for each disease) for the normal thyroid tissue samples from GTEx and fit a nonlinear curve with Locally Estimated Scatterplot Smoothing^45^ (LOESS) for each sex, using the score as a function of age. These sex-specific curves represented the dynamic trajectories of how TF-targeting patterns of these disease-related gene sets changed with age for males and females.

This produced two non-linear aging trajectories for each disease that capture how disease-associated gene regulatory patterns vary over lifetime in both males and females in ways that increase or decrease targeting of disease-related genes (**Figure 6**).

**Figure 6:**
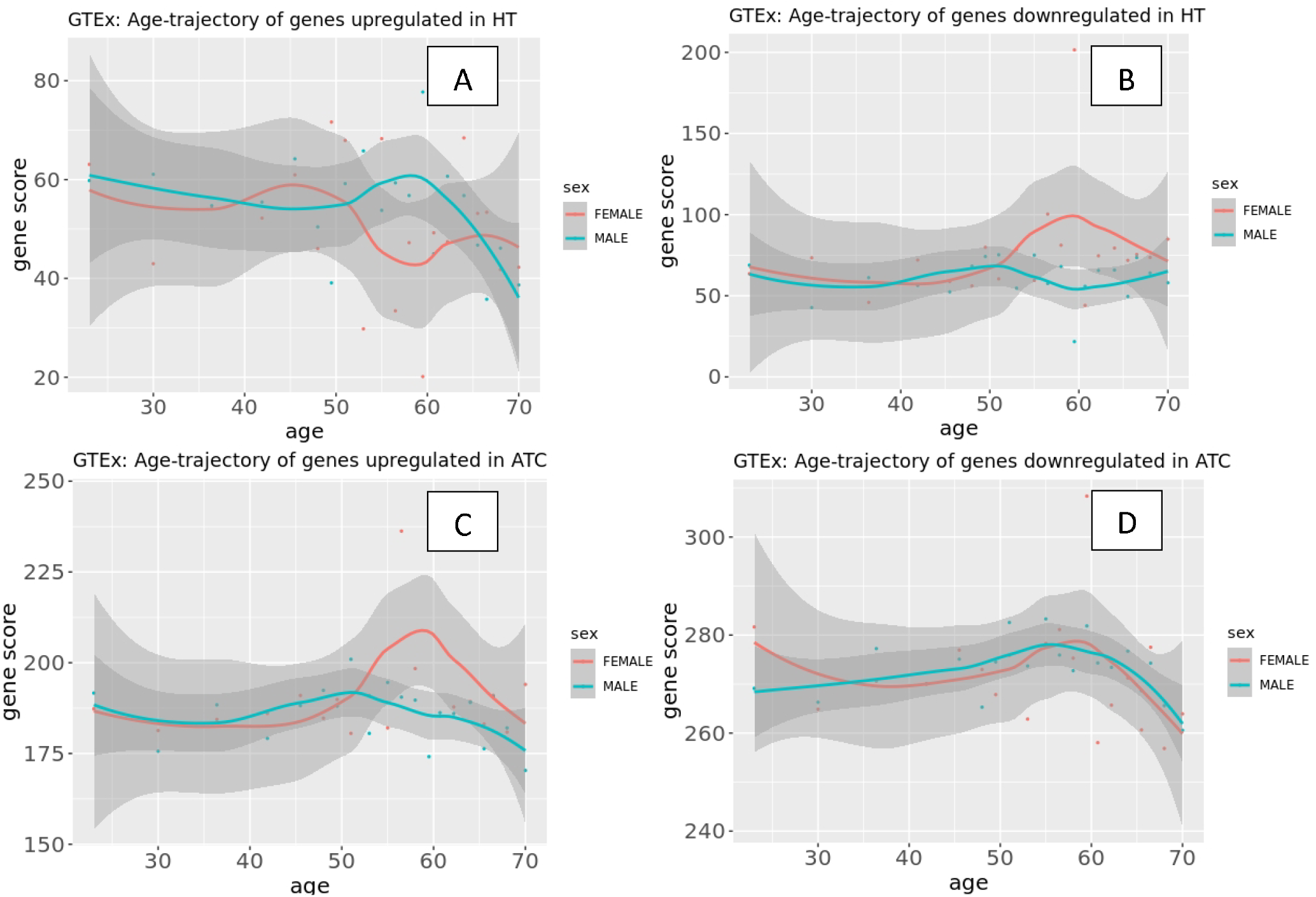
Sex-specific evolution of disease-associated genes with age in normal thyroid. The plot shows how gene targeting scores of disease-associated genes change with age for genes (A) targeted more in HT, (B) targeted less in HT, (C) targeted more in ATC and (D) targeted less in ATC, compared to normal thyroid. The x-axis represents age, and the y-axis represents gene targeting score of the corresponding gene set. Each plots shows an aging trajectory curve computed for male samples (colored teal) and a trajectory curve computed for female samples (colored reddish orange) in GTEx using Locally Estimated Scatterplot Smoothing (LOESS). The grey shaded region around each curve is the pointwise 95% confidence band.

Genes with greater targeting in HT compared to normal thyroid also had greater relative targeting in females than males between the ages of 40-50 and showed lower targeting in females in other age groups (**Figure 6 A**). The genes that had lower targeting scores in HT than in normal tissue, showed slightly lower targeting in females than males between ages 40-50, and higher targeting in females in other age groups (**Figure 6 B**). Thus, the pattern of female-specific TF-targeting between ages 40 and 50 was consistent with disease-related TF-targeting patterns in HT. This pattern aligns with the epidemiology of HT, which most commonly arises in middle-aged individuals and occurs more frequently in females than in males.

Genes that were less targeted in ATC than in normal thyroid, did not show prominent sex differences in TF-targeting as a function of age (**Figure 6 C**). However, genes that were highly targeted in ATC than in normal tissue, were more highly targeted in females than in males after age 50 and showed lower targeting in younger females (**Figure 6 D**). Thus, the female-specific TF-targeting in ATC is consistent with the directionality of disease-specific TF-targeting patterns after age 50, such that genes for which targeting increases in ATC are the same genes that are more highly targeted in females than males after age 50, consistent with epidemiological data on ATC, which is both more common in older adults above 50, and disproportionately affects females.

## Discussion

In this study, we used a novel computational framework to infer gene regulatory interactions at the individual level by combining BONOBO, a newly developed Bayesian model for estimating individual-specific co-expression networks, with PANDA, a message-passing algorithm that integrates these co-expression networks with prior knowledge on TF–gene motif and protein–protein interaction. We applied this framework to construct individual-specific gene regulatory networks from GTEx normal thyroid samples and identified genes whose targeting by TFs in the network models changed as a function of age in males and females. In normal thyroid, TF-targeting patterns of several proto-oncogenes, tumor suppressor genes and biological pathways involved in metabolism, immune response, cell adhesion and proliferation not only change with age in a non-linear way, but these aging-related non-linear patterns vary between males and females. Females undergo a marked shift in TF-regulatory patterns in the 40 to 60 age range, which is associated with similar changes in the targeting patterns by estrogen and androgen receptor TFs. Males exhibit milder inflection in TF targeting in the same age range, and do not show strong age-associated variation in sex hormone receptor TF targeting patterns. These results demonstrate the sex-specific roles of TFs, specifically estrogen and androgen receptor TFs, in the regulation of thyroid aging.

We compared gene regulatory networks between normal thyroid samples and those derived from individuals with HT and ATC. In both disorders, we found differential targeting of several pathways we had found to be aging-related, including those involved in immune response, metabolism and cell proliferation. However, we found differences between the two diseases in the patterns of altered regulation, compared to the age-regulated patterns in normal tissue. In HT, the directionality of disease-related regulatory changes was opposite to those observed in aging, but in ATC disease-related regulatory changes were in the same direction as aging-related changes. Further, in age groups where each disease is most prevalent (below 50 for HT and above 50 for ATC), TF-targeting profiles in females were more closely aligned with disease-related patterns, relative to what we saw in their male counterparts. These findings demonstrate that regulation of genes by TFs differ between the sexes in ways consistent with the appearance of HT at younger ages and a higher risk of ATC at older ages, while disproportionately affecting females more than males, at any age.

It is worth noting that even though our results show a strong association between the age-and sex-biased gene regulatory patterns and the prevalence of thyroid diseases, the gene regulatory networks were derived from observational data alone. Establishing a causal relationship between differential TF-driven gene regulation and disease risk would require experimental validation which is beyond the scope of this study. Another limitation of our study is the relatively small sample sizes in the HT and ATC datasets, which might reduce power of inference and hinder generalizability. However, the consistency of our findings across independent datasets provides some confidence in our conclusions. In addition, the normal thyroid population in GTEx primarily consisted of individuals of white and African American descent. The generalizability of our results to other ancestries may still be limited due to underrepresentation in our data. Further studies involving larger cohorts with diverse ancestry are necessary to confirm the validity of our results across the general population.

Furthermore, since GTEx does not include direct hormone measurements or menopausal status, the observed decline in sex hormone receptor targeting cannot be directly attributed to estrogen withdrawal, despite being consistent with the age-related decline in estradiol levels reported in NHANES. We therefore cannot determine whether the regulatory transitions identified here are a consequence of hormonal decline or reflect other age-related processes that co-occur during the same period.

The GEO datasets used for identifying gene regulatory changes in HT and ATC diseases, (GSE29315, GSE33630, GSE27155) did not include age and sex as clinical metadata. While we inferred biological sex from the genes on the Y chromosome and *XIST*, we could not determine age for individual samples. As a result, our conclusions regarding female disease-like regulatory patterns in specific age windows are based on indirect comparisons between the GTEx aging trajectories and the disease datasets, while adjusting for inferred biological sex. Prospective studies with complete clinical annotation, including both age and sex, would be needed to directly test whether the regulatory transitions identified here have causal implications for elevated disease risk in the relevant age groups.

A further limitation of our study is that we used bulk transcriptomic data for our analysis, which cannot fully distinguish changes in gene regulatory programs within a cell type from variability in cell state or cell composition proportions. Although we estimated immune cell proportions using ‘xCell’ and observed minimal age-associated changes, it is worth noting that subtle shifts in immune activation or stromal interactions could still influence the inferred network structure, particularly in aging tissues where functional reprogramming may occur independently of changes in cell abundance. Therefore, it is possible that some of the observed sex-specific regulatory changes may partly be driven by changes in cell state or cell composition. In future work, we plan to investigate single-cell or spatial transcriptomics approaches to identify these cell-type-specific gene regulatory changes.

Another point worth noting is that we have used LOESS smoothing to construct sex-specific smooth aging trajectories. Even though LOESS is a consistent estimator and our sensitivity analysis shows that our conclusions are robust with respect to parameter choices, subsampling and alternative smoothing methods such as the cubic spline, LOESS has some significant limitations including sensitivity to outliers and the tendency to miss subtle local variations. More sophisticated smoothing methods (such as generalized additive models with explicit nonlinearity inference) on larger datasets with more even age distributions is required to establish the generalizability of our findings regarding the molecular changes in females between ages 40 and 60.

Despite these limitations, the analyses presented here provide new and powerful insight into how age and sex influence gene regulatory processes and shape thyroid health. We found that sex hormone receptor transcription factors, particularly estrogen and androgen receptors, play a central role in mediating non-linear, age-dependent and sex-specific shifts in gene regulation. The patterns we found are consistent with sex disparities in thyroid disease risk and with evidence of biases in the age of onset, particularly in women. Notably, we found that there are specific sex- and age-related windows in which gene regulatory changes are most significant and correlate with disease onset, suggesting that these may be useful in finding potential targets for intervention to reduce thyroid disease burden and improve patient outcomes.

## Data and Code Availability

Raw data to construct gene regulatory networks and other analysis were downloaded from open-source databases dbGap^47^, Recount3^48^, GEO^21^, STRINGdb^49^, CIS-BP^50^ and NHANES^24^. The sex-specific TF-gene and PPI prior networks are publicly available through the GRAND^51^ database.

Results from gene set enrichment analysis for all datasets are available as supplementary tables.

Sample-specific gene regulatory networks are stored in an Amazon Web Services S3 bucket and will be made publicly available upon acceptance.

List of aging-associated and disease-associated genes are available as supplementary data.

R codes for downstream analysis are available on a GitHub public repository: https://github.com/Enakshi-Saha/Aging-Thyroid

## Acknowledgements

This work was supported by grants from the University of South Carolina Office for the Study of Aging (OSA). **ES** was supported by OSA research fellowship in aging.

## Author Contributions

**ES:** Conceptualization, Data curation, Formal analysis, Investigation, Methodology, Software, Validation, Visualization, Funding Acquisition, Writing – original draft.

## Methods

### RNA-sequencing Data

We downloaded RNA-Sequencing data for 706 normal thyroid tissue samples from the GTEx Project from the Recount3 database^48^ using R package ``recount3” (version 1.4.0). Clinical data for these samples were obtained from dbGap (https://dbgap.ncbi.nlm.nih.gov/) under accession number phs000424.v8.p2. From these, we removed 53 samples that the GTEx consortium had identified as “biological outliers” for various reasons (as described in https://gtexportal.org/home/faq). We performed our final analysis on the remaining 653 samples (434 males and 219 females). **Table 1 shows** the distribution of clinical variables in GTEx by sex.

**Table 1:** Clinical characteristics of the GTEx data by sex.

|  | Female | Male |
| --- | --- | --- |
| **Sample size** | 219 (33.53%) | 434 (66.46%) |
| **Age** |  |  |
| *Mean ± std (range)* | 52.15 ± 12.64 (21-70) | 53.62 ± 12.48 (20-70) |
| **Race** |  |  |
| *White (%)* | 186 (84.93%) | 372 (85.71%) |
| *Black or African American (%)* | 27 (12.33%) | 48 (11.06%) |
| *Others (%)* | 6 (2.74%) | 14 (3.22%) |
| **BMI** |  |  |
| *Mean ± std (range)* | 26.97 ± 4.16<br>(18.58-35.42) | 27.86 ± 3.96<br>(18.64-35.05) |
| **RIN** |  |  |
| *Mean ± std (range)* | 6.80 ± 0.73<br>(5.5-9.7) | 6.74 ± 0.63<br>(5.5-9.3) |
| **Smoking status** |  |  |
| *History of smoking (%)* | 140 (63.93%) | 304 (70.04%) |
| *No history of smoking (%)* | 73 (33.33%) | 125 (28.8%) |
| *NA (%)* | 6 (2.74%) | 5 (1.15%) |
| **Ischemic time (hours)** |  |  |
| *Mean ± std (range)* | 6.75 ± 6.96<br>(0.0-24.4) | 8.5 ± 7.11<br>(0.0-27.45) |

RNA-sequencing data were normalized using the Transcripts per Million (TPM) method, by the ``getTPM” function in the Bioconductor package “recount” (version 1.20.0)^52^ in R (version 4.1.2). We filtered out genes that were lowly expressed, keeping only 26792 genes (including 64 Y chromosome genes) which had transcript counts of at least 1 TPM in at least 10% of samples. To build gene regulatory networks, we further filtered out genes that were not present in the TF/target motif prior, leaving 26507 genes to be used in gene regulatory network inference and subsequent analysis.

To validate the aging-related genes and pathways we discovered using data from GTEx, we used a RNA-sequencing dataset from the Gene Expression Omnibus (GEO)^21^, with accession number GSE165724^53^. This dataset contained expression of 63299 transcripts (including 554 transcripts on the Y chromosome) from normal thyroid samples from 12 individuals without any thyroid disorders. We normalize gene expression using TPM, which requires dividing the transcript counts by the gene length and so we removed genes for which base-pair lengths were unavailable, leaving 61716 transcripts. For genes with multiple transcripts, we kept only the transcript with the highest variability across samples. Finally, we removed genes with counts < 1 TPM in at least 10% samples, just as we did in GTEx. This left 25862 genes. To build gene regulatory networks, we further filtered out genes that were not present in the motif prior, leaving 18912 genes for our final analysis.

Samples in GSE165724 were all males with age ranging between 48 and 79 (with mean 60.33 and standard deviation 9.2278). No other clinical variables were available for this dataset.

We log transformed the TPM normalized gene expression data to be used as input for computing individual-specific TF-gene regulatory networks.

To verify whether the self-reported gender in GTEx and GSE165724 match with the chromosomal complement biological sex, we performed a principal component analysis on the expression levels of genes on the Y chromosome. We found that the self-reported gender perfectly aligned with the biological sex defined by the presence (for males) and absence (for females) of the Y chromosome (**Figure S9**). We also found that due to mis-mapping of transcripts, some genes on the Y chromosome were assigned non-zero expression in females due to mis-mapping of transcripts. For these genes, we set expression to zero for biological females.

### Microarray Data

We used three microarray gene expression datasets from the Gene Expression Omnibus (GEO)^21^ (accession numbers GSE29315, GSE33630^54^ and GSE27155^55,56^). We used GSE29315 (6 samples with HT and 8 control samples) was used as the validation dataset for identifying HT-related gene regulatory changes. For identifying ATC-related gene regulatory changes, we used GSE33630 (11 samples with ATC and 45 control samples) and GSE27155 (4 samples with ATC and 4 control samples), as discovery and validation datasets respectively. These datasets did not have any clinical data available other than disease status.

For GSE29315, mRNA assays were performed using Affymetrix Human Genome U95 Version 2 Array and expression values were recorded for 12625 probes. Data were then normalized with the Robust Multiarray Average (RMA) method, using the affy Bioconductor package^57^. For GSE33630, mRNA assays were performed using Affymetrix U133 Plus 2.0 arrays and expression were recorded for 54675 probes. Expression data were processed using the quantile-normalized trimmed-mean method. Data were normalized with RMA, using the affy package. For GSE27155, mRNA assays were performed using Affymetrix HG_U133A arrays and expression values were recorded for 22283 probes. Data was processed using the Ann Arbor quantile-normalized trimmed-mean method.

In all microarray datasets, for the genes with multiple transcripts, we kept only transcripts with the highest variability. We further filtered out genes which were not present in the TF motif prior, thus leaving 9173, 23349 and 13436 genes for GSE29315, GSE33630 and GSE27155, respectively. Normalized expression data were log2-transformed before constructing the individual-specific gene regulatory networks.

None of these datasets included any clinical variables (e.g. age and sex) as metadata and hence we could not provide a summary table as we did in **Table 1 for** GTEx.

### Determining Biological Sex Using Expression Levels of Y Chromosome Genes and XIST

For all GEO datasets for which sex information was unavailable (GSE29315, GSE33630, GSE27155), we inferred biological sex by performing principal component analysis (PCA) on the expression levels of Y chromosome genes. In all three datasets, samples clustered into two well-separated groups along PC1, which were consistent with the difference in *XIST* expression levels across the two clusters. We annotated samples with high *XIST* expression as females and samples with low *XIST* expression as males (**Figure S10**). We used the inferred sex to assign sex-specific motif and PPI priors while fitting individual-specific gene regulatory networks. We also used the inferred sex to adjust for sex effects in our ‘limma’ models while determining disease-specific gene regulatory interactions. It is worth noting that each of the datasets contained normal thyroid samples and samples from multiple diseases. We used all samples for sex identification to ensure robust cluster separation, but included only normal thyroid samples, Hashimoto’s Thyroiditis (HT) and Anaplastic Thyroid Carcinoma (ATC) in the study for the final analysis.

### Estimating Individual-specific Gene Regulatory Networks

We used normalized and log transformed gene expression data from the discovery and validation datasets for both normal thyroid and our disease models to compute individual-specific gene-gene co-expression networks using an empirical Bayesian model BONOBO^22^, which integrates individual-specific expression data with a population-specific prior co-expression network to infer a posterior gene-gene co-expression network for each individual, thereby borrowing strength across samples. The adjacency matrices of the co-expression networks estimated by BONOBO were then used as inputs to PANDA^23^, a message passing algorithm for inferring individual-specific TF-gene regulatory networks.

PANDA reconstructs gene regulatory networks by integrating multiple sources of omics data—typically a motif prior W (transcription factor–target gene potential interactions), a protein–protein interaction (PPI) network P among transcription factors, and a gene co-expression network C among target genes. PANDA interprets each as describing relationships among entities that share information that reflects the regulatory process such that TF–TF similarities (from P) should correspond to similar regulatory targets in W; gene–gene co-expression (from C) should correspond to similar upstream regulators in W. It does this through an iterative message passing algorithm that seeks self-consistency among corresponding three network matrices, W, P, and C, updating them at each stage in the iterative process until the process converges, producing a final regulatory network model.

Because we wanted individual-specific regulatory network models for both sexes, as inputs to PANDA, we used the individual-specific co-expression adjacency matrices coming from BONOBO, together with the appropriate sex-specific TF-gene motif regulatory network priors (male or female as defined by allosomes content, as defined below), and TF-TF PPI priors. PANDA was implemented in python using package netZooPy (version 0.9.10)^58^. This produced a collection of individual-specific gene regulatory networks specific to each individual’s pattern of expression and adjusted for their sex.

For each of the resulting individual-specific gene regulatory networks, we computed summary measures that included gene in-degree, or “gene targeting score,” equal to the sum of all incoming edge weights from all TFs to that gene. Similarly, for each TF, we calculated the TF out-degrees, or “TF targeting score,” equal to the sum of all out-going edge weights from a specific TF to its target genes. Finally, because we were interested in understanding the cumulative effects of regulatory changes on biological processes, we calculated “gene set scores” or “pathways score” defined as the sum of all in-coming edges from TFs to all genes in a specific gene set or biological pathway. For clarity, the definitions of these summary measures are provided in **Table 2**.

**Table 2:** Definitions of network summary statistics used in the analysis.

| Term | Definition |
| --- | --- |
| Gene Score or Gene In-degree | Sum of incoming edge weights from all TFs to the specific gene |
| TF Score or TF Out-degree | Sum of out-going edge weights from the specific TF to all its target genes |
| Pathway Targeting Score | Sum of incoming edge weights from TFs to all the genes in the specific pathway |

The prior networks required for PANDA were constructed as follows.

### Sex-specific Transcription Factor-Gene Motif Prior

The TF-gene motif prior is a bipartite network where the network edges connect TFs to their target genes, such that the edge value between a TF and a gene is either 1 or 0, depending on whether that TF motif is present or absent within the promoter region of that gene. To create this prior motif network, we downloaded TF motif position weight matrices (PWM) for *Homo sapiens* with direct or inferred evidence from the Catalog of Inferred Sequence Binding Preferences (CIS-BP) Build 2.0^50^. These PWMs were mapped to the human genome (hg38) using FIMO^59^. We kept only the highly significant matches (p-value < 10e-5) within the promoter regions of Ensembl genes (for which GENCODE v39 annotations were obtained from http://genome.ucsc.edu/cgi-bin/hgTables). We defined promoter regions as the interval of [-750; +250] base pairs centered around the transcription start site. The result was a prior motif network with binary regulatory edges for 997 TFs, that collectively targeted a total of 61,485 genes.

To enable statistical comparisons between networks from males and females, we needed to ensure that the prior motif networks for both sexes had identical sets of edges. To account for the absence of Y chromosome genes in females, we defined sex-specific TF-gene regulatory priors as follows. To construct the female-specific regulatory prior, we assigned an edge weight of zero to the 52,266 edges originating from, or connecting to, Y chromosome TFs or genes.

### Protein-protein Interaction Prior

The PPI interaction network connects two TF proteins with an edge if these two proteins are evidenced to chemically interact with each other. To construct this network, we first downloaded PPI data from the String database^49^ (version 11.5) using the STRINGdb Bioconductor package. We then filtered the PPI data, using a score threshold index of 0, keeping only interactions between the TFs that were present in the TF-motif network. To make the PPI scores consistent, we normalized them by dividing each score by 1000, thus restricting the values to a uniform range of 0 to 1 for both the PPI network and the TF-motif network. For each TF, we set the self-interaction value to one. Finally, we transformed the adjacency matrix into a symmetric PPI matrix as PPI networks are essentially undirected.

### Differential Targeting Analysis

Our goal in this analysis was to find regulatory changes that occur with age, of those that differentiate between the sexes and changes associated with disease. Because we had multiple datasets and different parameter types—age, a continuous variable, and sex and disease status, both discrete variables—we needed appropriate models to find differentially interacting TFs, genes, and pathways.

#### Model 1-age

To identify the TF-driven gene regulatory changes associated with aging in GTEx (excluding individuals with HT) and the independent validation dataset GSE165724, the targeting scores for each gene were regressed on age, using a gene-specific linear regression model. These linear models were fit using the R package “limma” (version 3.50.3)^60^, in which we used empirical Bayes to stabilize variance and then calculated p-values for the model coefficients using Student’s t-test. False discovery rate (FDR) was corrected using the Benjamini Hochberg^61^ procedure with significance cutoff set at 0.05. For GTEx, the linear models accounted for the effects of age, while adjusting for the effects of relevant confounders, such as sex (male and female), race (White, Black or African American, Others, and Unknown), BMI, smoking status (ever-smoker and never-smoker), sample ischemic time, RNA integrity number (RIN), and batch. For the GSE165724 dataset, no other clinical variables other than age and sex were available and so these were the only variables used in the model.

#### Model 2-sex

To identify the sex-specific regulatory changes associated with aging in GTEx, the gene targeting scores for each gene were regressed on sex, age and the *interactions between age and sex*, using a gene-specific linear regression model. These linear models were fit using the R package ``limma” and adjusted for the effects of confounders, such as race, BMI, smoking status, ischemic time, RIN and batch.

#### Model 3-disease status

To estimate regulatory changes associated with HT in GTEx, the gene targeting scores for each gene were compared between networks constructed from individuals with and without HT, using linear regression models, fitted with R package ``limma”. The linear models accounted for the effects of disease status (with and without disease), while adjusting for the effects of age, sex, race, BMI, smoking status, ischemic time, RIN and batch. To estimate the regulatory changes associated with ATC in GSE33630 and to validate the findings on HT and ATC using the validation datasets GSE29315 and GSE27155, respectively, linear models were used with only disease status as covariate, as no other clinical data were available.

### Gene Set Enrichment Analysis

We performed ranked Gene Set Enrichment Analysis (GSEA) using R package ``fgsea”^62^ (version 1.20.0) and gene sets from the Kyoto Encyclopedia of Genes and Genomes (KEGG), as downloaded from the Molecular Signatures Database (MSigDB)^63^. After filtering out genes that were not present in the expression datasets, we only considered gene sets containing between 15 and 500 genes. Genes were then ranked by the t-statistics obtained from the limma differential targeting analysis and the ranked list was used as input for the ranked GSEA. We corrected for FDR using the Benjamini-Hochberg procedure.

### Cell Type Deconvolution Analysis

We performed cell type deconvolution analysis on the normal thyroid samples from GTEx to understand how the proportion of different cell types varies with age in normal thyroid. We used “xcell”^34^ on the unfiltered GTEx gene expression data with R package “immunedeconv” (version 2.1.0) to estimate the composition of immune and stromal cells. For each cell type, we fit a linear model (using R function “lm”) to predict the proportion of the corresponding cell type as a function of age, while adjusting for clinical covariates such as sex, race, BMI, smoking status, ischemic time, RIN and batch, enabling us to quantify how cell type composition in normal thyroid varies with age.

### Constructing Gene Set and TF-specific Aging Trajectories for Normal Thyroid Samples from GTEx

We divided the aging-related pathways identified in the differential targeting analysis of the GTEx samples into six broader categories based on their functions, following the KEGG hierarchy of pathways. These are (i) Immune system and immune system diseases, (ii) Cell signaling, cell growth and death, (iii) Cell communication, (iv) Genetic Information processing (transcription, translation, replication & repair), (v) Endocrine system, circulatory system, Nervous system and related diseases and (vi) Metabolism (carbohydrate and lipid). For each of these categories of pathways, we defined a gene-set-specific gene targeting score for every individual, defined as the sum of incoming edge weights from TFs to all the genes in that set within the gene regulatory network for that individual. Then we divided samples for each sex into 20 consecutive age-groups by age quantiles in the GTEx. Then within each age group and each sex, we computed the medians of the gene targeting scores across individuals in that age and sex group. For each pathway category, a scatterplot of the gene targeting scores (y-axis) over the mid-point of each age-group (x-axis) was drawn and colored by sex (reddish orange for females and teal for males). Next, for each gene set (i.e. pathway group), we fit a nonlinear curve for each sex with LOESS, using the targeting score for that gene set as a function of age, with R package ggplot2. These sex-specific curves represent the trajectories of how the TF-targeting patterns of these gene sets vary with age for males and females. Thus, for each sex, we get a non-linear aging trajectory for each gene set, which demonstrates how gene regulatory patterns vary with age to increase or decrease targeting of genes in that specific group of pathways, specific to each sex.

For estrogen and androgen receptor TFs, ESR1, ESR2 and AR, we computed the aging-gene-specific out-degree of the TF for each individual subject, defined as the sum of all outgoing edge weights from that TF to all the aging genes within the gene regulatory network for that individual. For each TF we then fit a nonlinear curve with LOESS (using ggplot2) for each sex, using the TF out-degree as a function of age and stratifying the samples into age and sex groups following the procedure mentioned above. These sex-specific curves represent the dynamic trajectories of how the estrogen and androgen receptor TFs target genes over varying age in each sex. Thus, we get non-linear aging trajectories for ESR1, ESR2 and AR for each sex.

We demonstrate the robustness of our trajectory inference approach in the following section (**Figure S4** - **Figure S7**). In addition, we demonstrate that fitting LOESS on raw ages without aggregation within age groups (bins) produces largely flat curves for both sexes (**Figure S8**), as sample-level variability at each age obscures the underlying age-related trends. Aggregating samples into age groups before fitting (**Figure 3**) reduces this noise and reveals the sex-specific aging trajectories, including the transitions between ages 40 and 60 observed in females. Aggregating samples into bins before applying smoothing curves is a standard practice for visualizing large and noisy data for ensuring scalability and noise removal^64,65^.

### Sensitivity Analysis on the Gene Set-specific Aging Trajectories for Normal Thyroid Samples from GTEx

To assess the robustness of the non-linear aging trajectories observed in GTEx, we performed four sensitivity analyses on the LOESS-smoothed pathway targeting score trajectories described above. In all analyses, pathway targeting scores were computed as the mean in-degree across genes belonging to each pathway superclass, aggregated by age group and sex prior to smoothing, as in the main analysis.

First, we examined sensitivity to the choice of smoothing parameter by fitting LOESS curves with three alternative span values (0.6, 0.75, and 0.9) and overlaying them on the same plot (**Figure S4**). To quantify uncertainty around the default span (0.75), we additionally computed pointwise 95% bootstrap confidence intervals. We note that the trajectories were well-aligned irrespective of the LOESS smoothing parameter and retained the inflection points around ages 40 and 60 for females, as observed in the main analysis.

Second, we assessed whether the observed transitions could be driven by any specific subset of samples or by uneven sample density across age groups (**Figure S5**). We repeatedly fitted LOESS curves (span 0.75) on 100 random subsamples, consisting of 80% of individuals from the full data, drawn without replacement, aggregating by age group before smoothing in each iteration. We overlaid these resulting 100 curves on the same figure as our original trajectories drawn using the full sample.

Third, we compared the LOESS trajectories to natural cubic spline regression fits as an alternative modeling approach (**Figure S6**). We placed spline knots at the 25th, 50th, and 75th percentiles of the observed age distribution in GTEx. Even though the knot placements were entirely determined by the data without reference to the inflection points observed at ages 40 and 60 for females, we still observed that the trajectories formed by the cubic splines and the original LOESS trajectories from the main analysis were well-aligned, providing independent support for the observed patterns, rather than a consequence of modeling assumptions.

Fourth, we fit a LOESS trajectory after leaving out one of the 20 bins used in the original analysis (**Figure S7**), drawing 20 such trajectories to ensure that the observed patterns in females around ages 40 and 60 are not driven by samples in any specific age group. We observe that while the degree of curvature varies depending on which age group is excluded, the change in female trajectory between ages 40 and 60 persists across all 20 leave-one-bin-out trajectories.

## Supplementary Material

**Figure S1:**
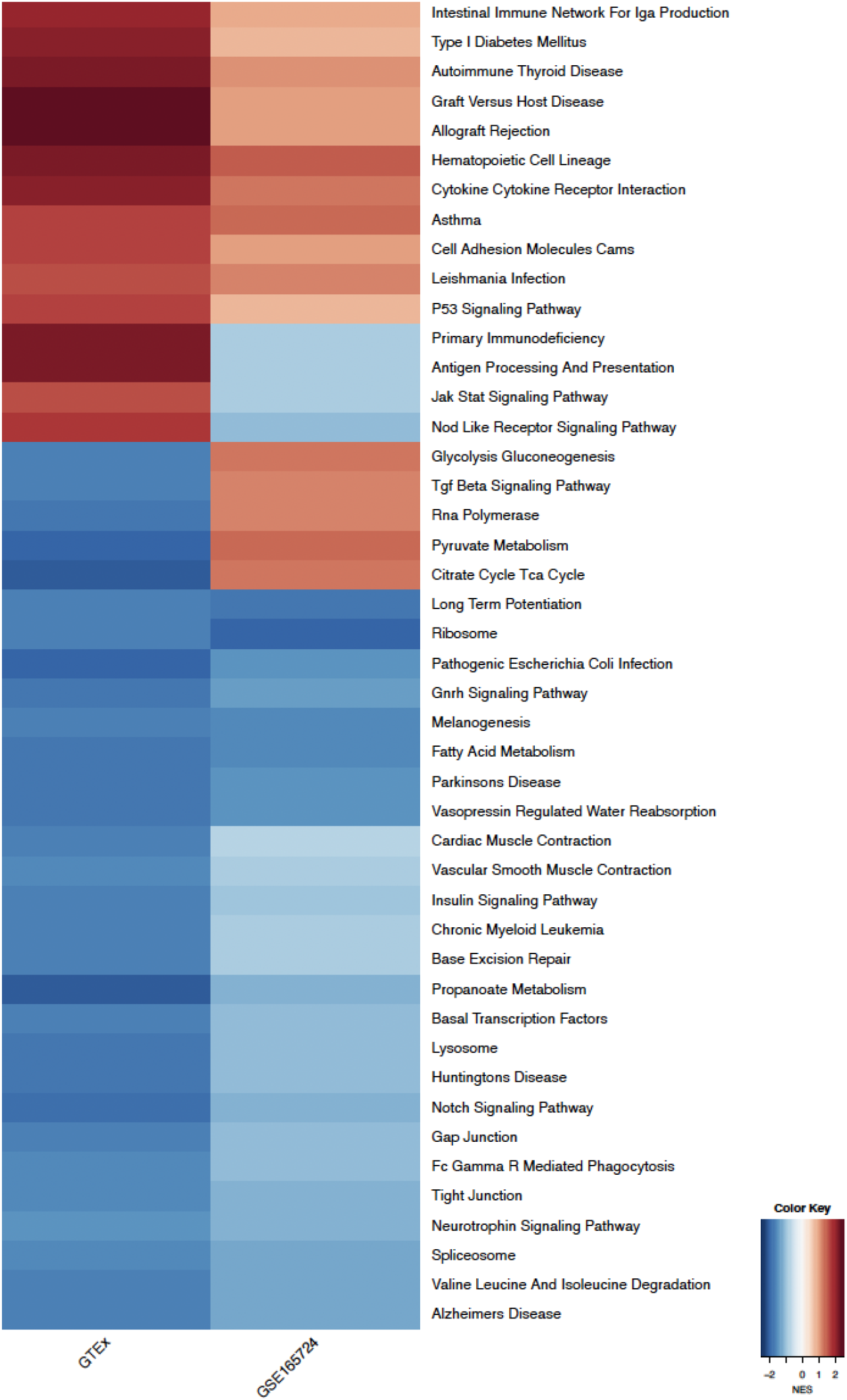
Biological pathways significantly differentially targeted by transcription factors (TFs) with age. The heatmap shows the normalized enrichment scores (NES) of aging-associated pathways identified b gene set enrichment analysis (GSEA) in GTEx. Rows represent pathways. The first column shows NES for GSEA in GTEx, and the second column shows NES for GSEA in the validation dataset GSE165724. Pathways marked red and blue are increasingly and decreasingly targeted by TFs with age, respectively.

**Figure S2:**
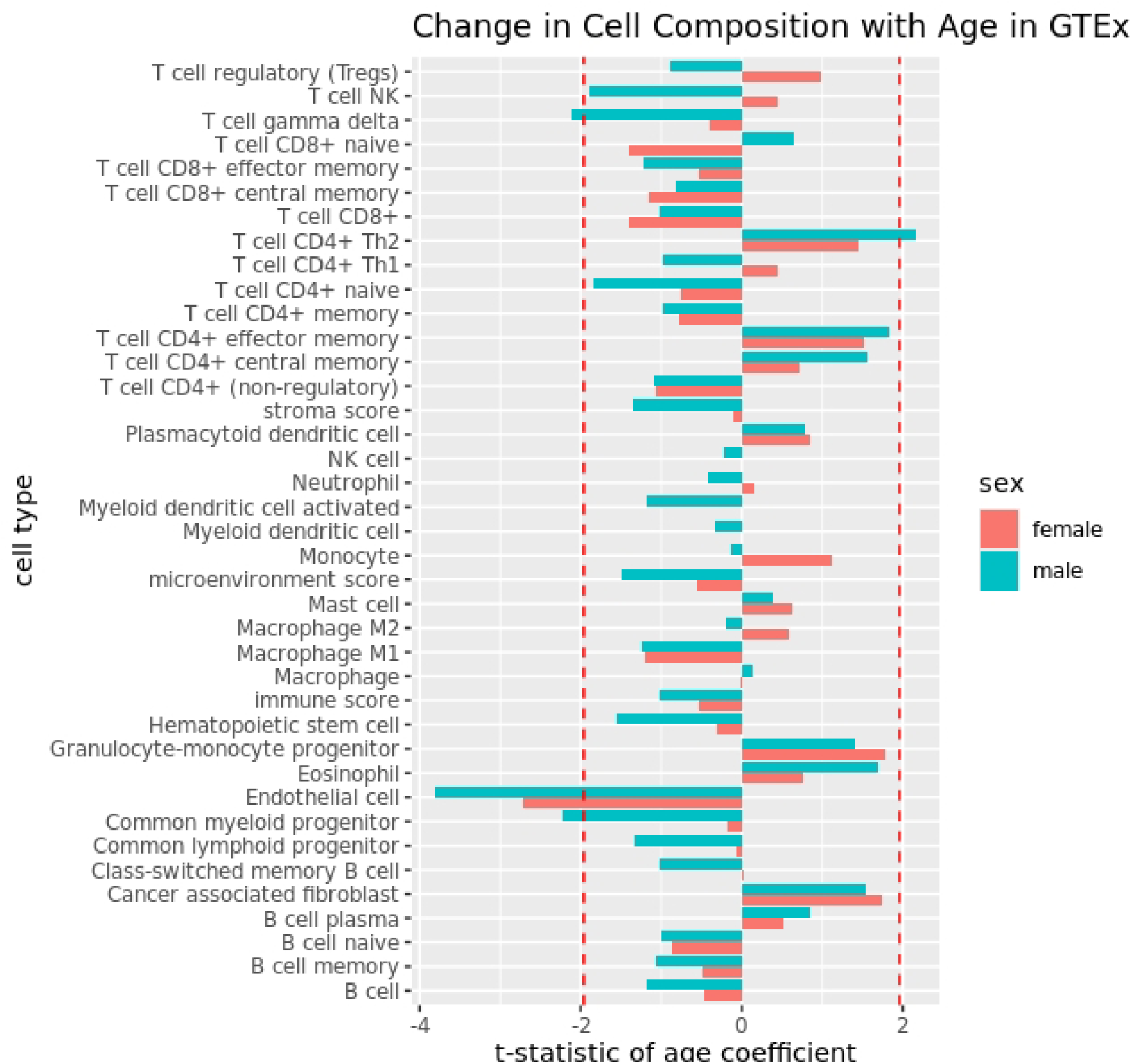
Change in immune and stromal cell composition with age in GTEx thyroid samples: for each cell type, the bar lengths correspond to the t-statistics of the age coefficients from linear models with cell type proportion as response and age as covariate, while adjusting for other clinical covariates. Vertical red dotted lines show the 2.5% and 97.5% quantiles of the standard normal distribution. Cell types for which the corresponding bars cross these lines are inferred to be significantly (p-value < 0.05) changing in proportion with age in normal thyroid.

**Figure S3:**
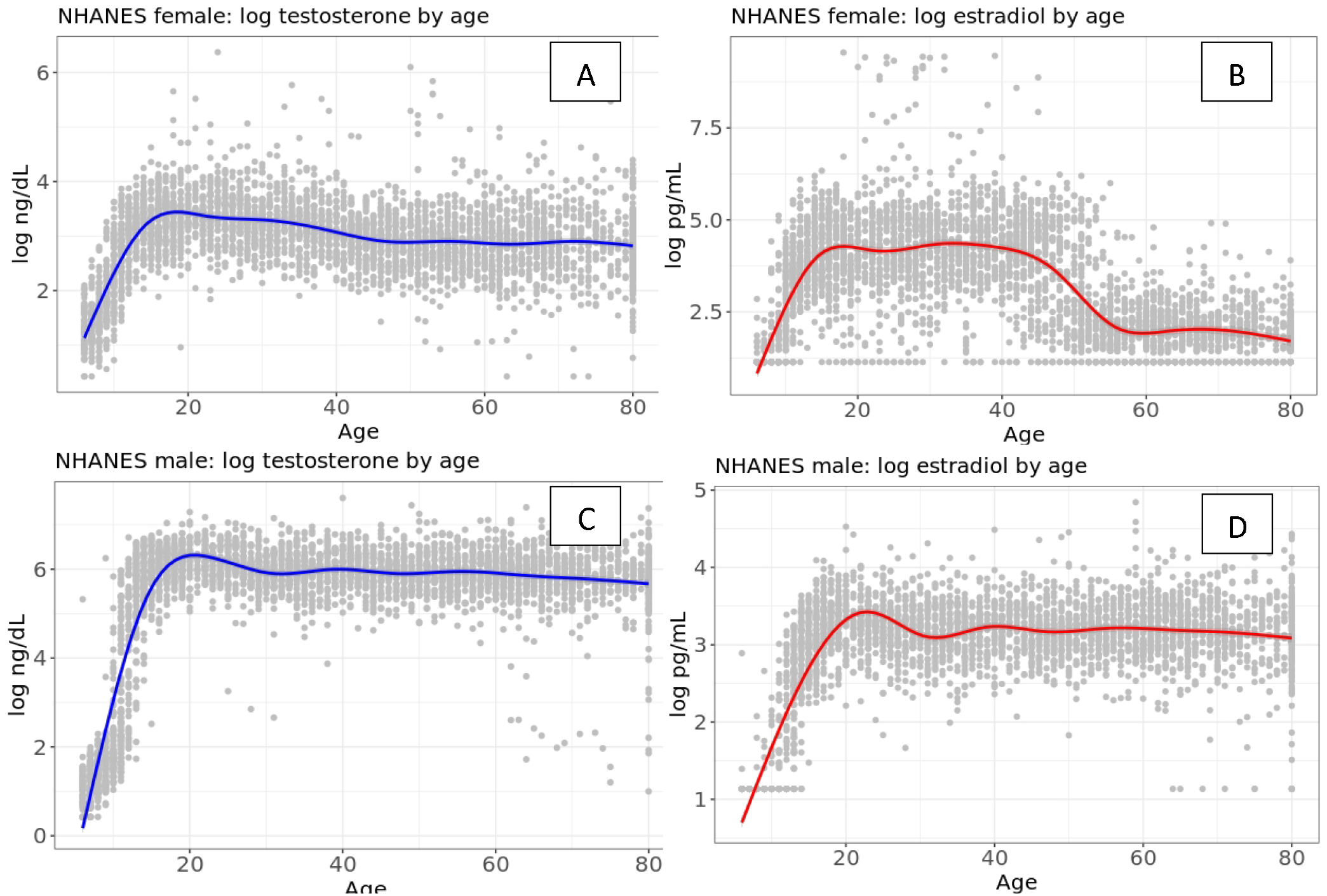
Change in sex steroid hormone levels with age in individuals in the National Health and Nutrition Examination Survey (NHANES) 2015-16. The plot shows logarithm of the levels of (A) testosterone in females, (B) estradiol in females, (C) testosterone in males and (D) estradiol in males. The red and blue smooth curves are drawn using Locally Estimated Scatterplot Smoothing (LOESS) for testosterone and estradiol respectively.

**Figure S4:**
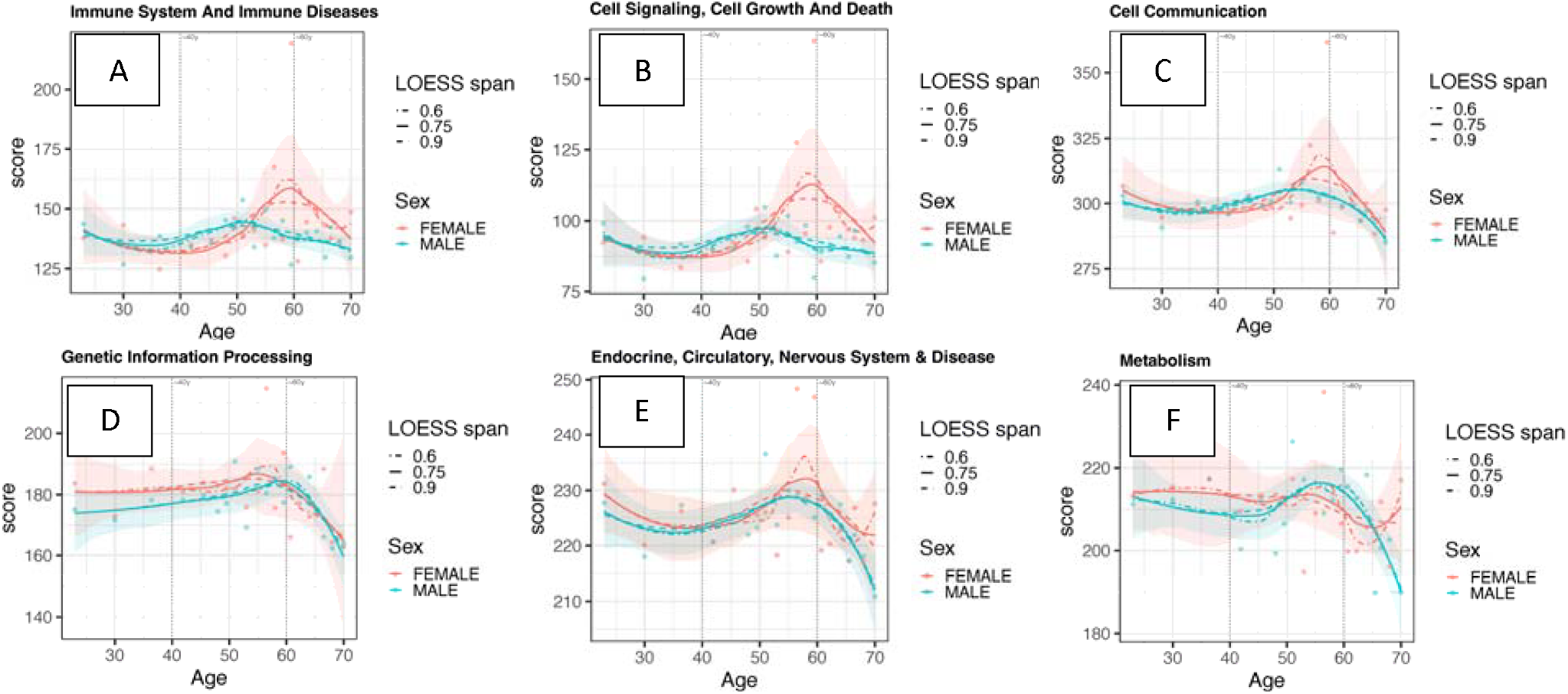
Robustness of sex-specific aging trajectory curves to the choice of LOESS smoothing parameter. Each plot shows the aging trajectories for male (teal) and female (reddish orange) samples in GTEx for a group of biological pathways (grouped according to KEGG hierarchy). The pathway targeting score on the y-axis is the mean in-degree of all genes in that group of pathways, aggregated across samples within each age group. The x-axis represents age. Three LOESS smoothing spans are overlaid per plot: 0.6 (dot-dash), 0.75 (solid, the default used in the main Figure 3), and 0.9 (dashed). The shaded ribbon represents the 95% point-wise confidence intervals around the default span (0.75). Vertical dotted lines mark ages 40 and 60 years. Consistent inflection patterns across all three spans indicate that the observed non-linear transitions in females are not an artifact of the chosen smoothing parameter.

**Figure S5:**
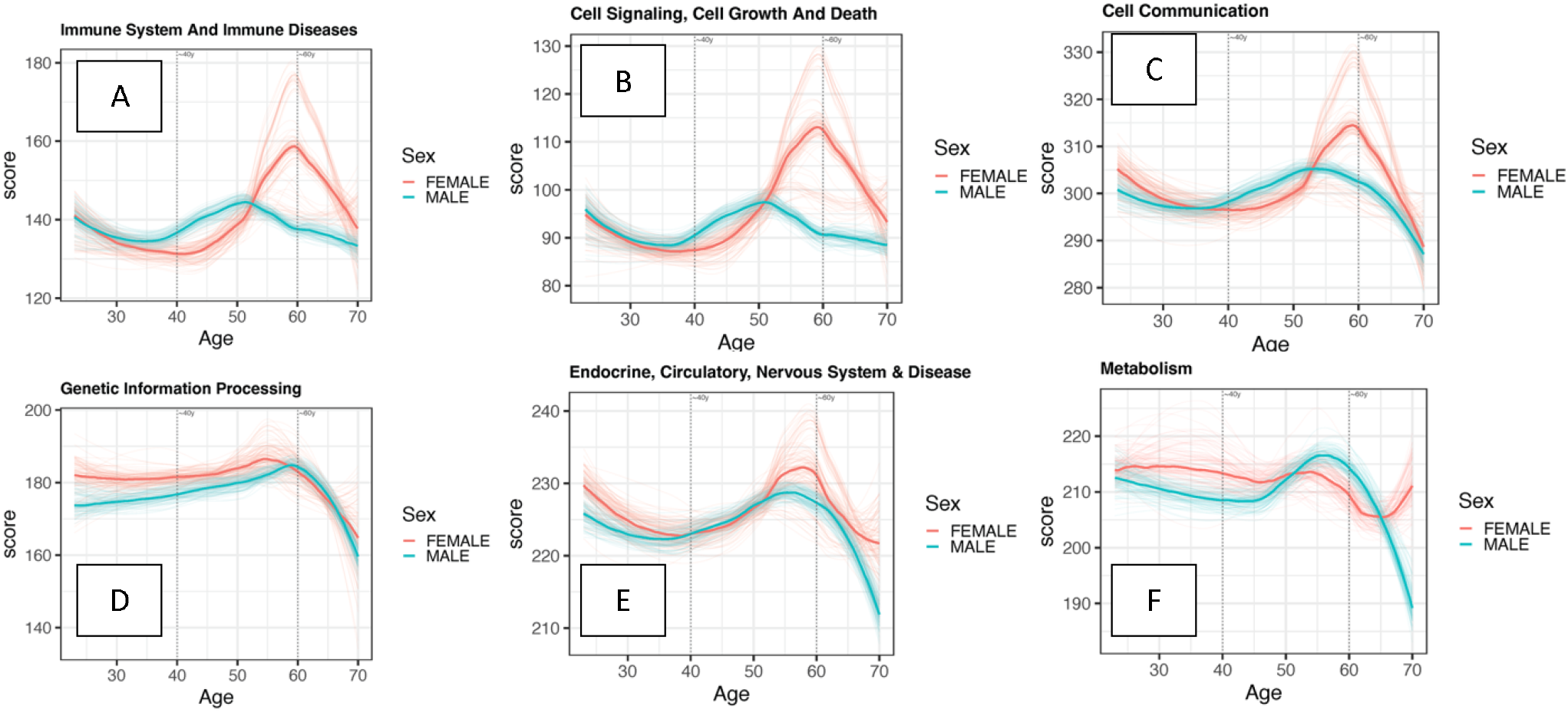
Robustness of sex-specific aging trajectory curves to the composition of the sample. Each plot shows the aging trajectories for male (teal) and female (reddish orange) samples in GTEx for a group of biological pathways (grouped according to KEGG hierarchy). The pathway targeting score on the y-axis is the mean in-degree of all genes in that group of pathways, aggregated across samples within each age group. The x-axis represents age. Each plot overlays 100 LOESS trajectories (faint lines), each fitted on a random 80% subsample of individuals without replacement, aggregated by age group prior to smoothing. The bold line represents the median trajectory across all 100 subsampled runs. Vertical dotted lines mark ages 40 and 60 years. Consistent trajectory shapes across all subsamples indicate that the inferred non-linear transitions are not driven by any subset of samples or by uneven sample density across age groups.

**Figure S6:**
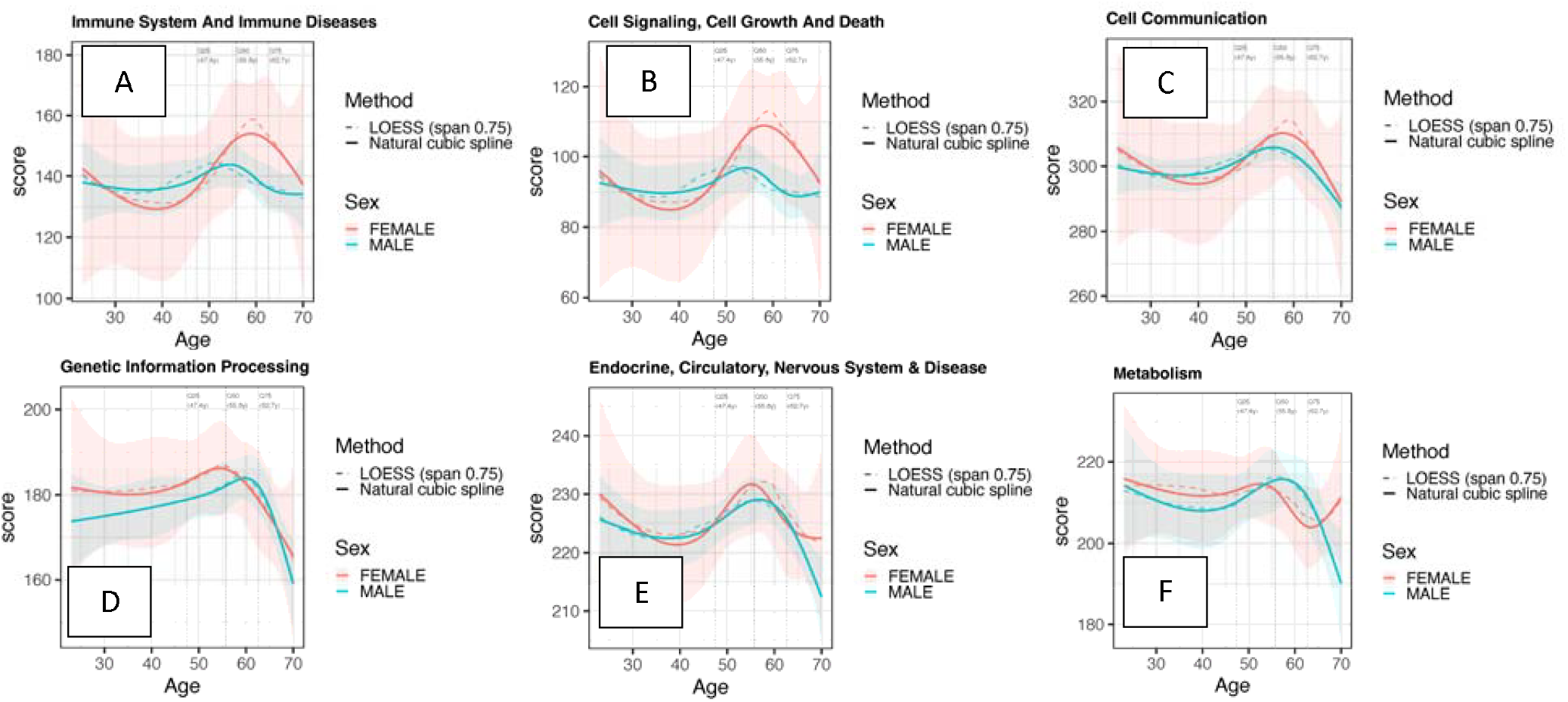
Robustness of sex-specific aging trajectory curves to the choice of smoothing method. Each plot shows the aging trajectories for male (teal) and female (reddish orange) samples in GTEx for a group of biological pathways (grouped according to KEGG hierarchy). The pathway targeting score on the y-axis is the mean in-degree of all genes in that group of pathways, aggregated across samples within each age group. The x-axis represents age. Each plot overlays a natural cubic spline regression (solid line, ±95% confidence interval ribbon) and a LOESS curve with span 0.75 (dashed line, the default used in the main Figure 3). Spline knots were placed at the 25th, 50th, and 75th percentiles of the observed age distribution, purely data-driven and without reference to ages 40 or 60; vertical dotted lines mark where these knots fell. Since the knot locations were determined by the data rather than pre-specified at ages 40 or 60, the agreement between the spline and LOESS trajectories suggests that the observed transitions are not an artifact of the smoothing method.

**Figure S7:**
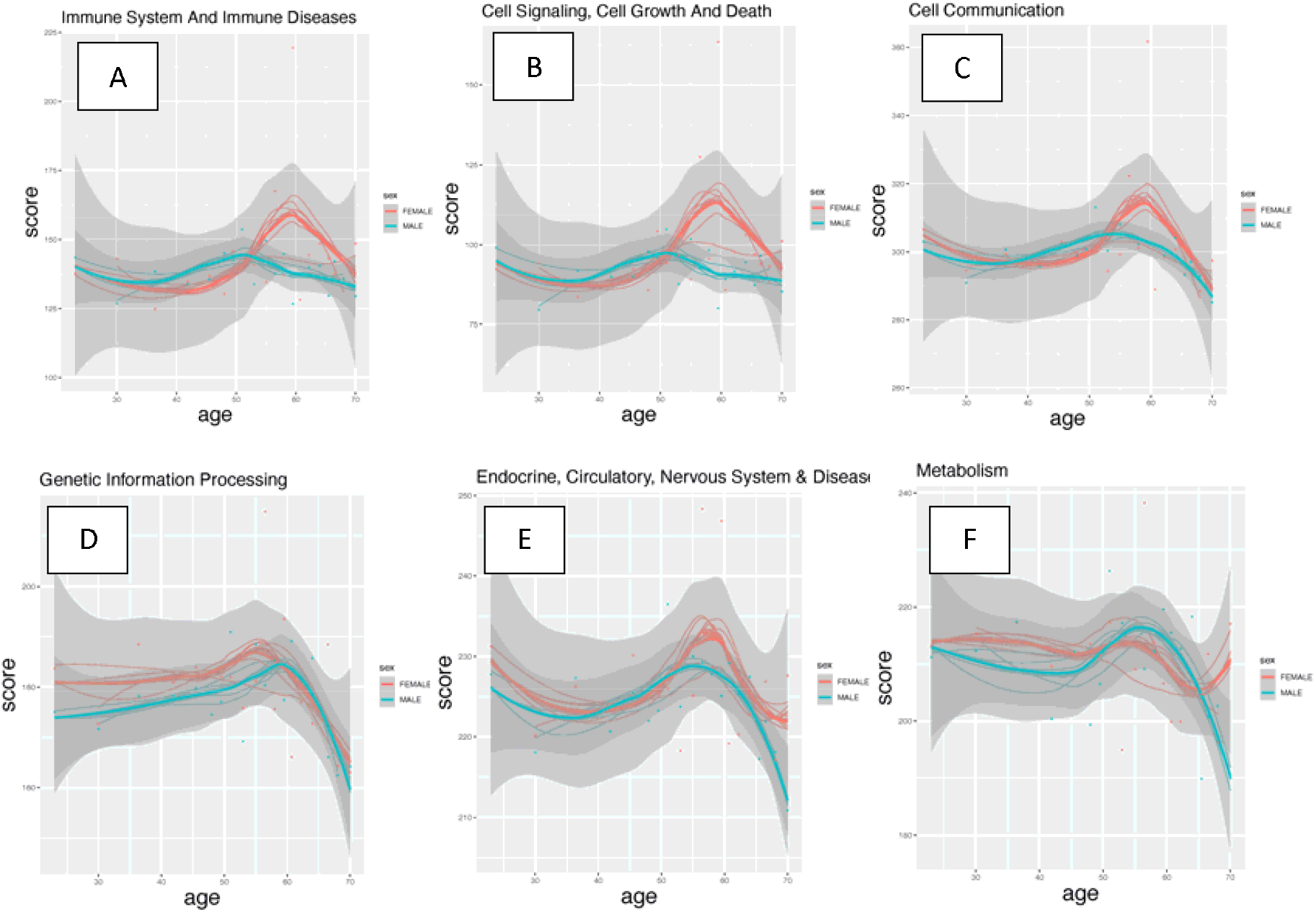
Robustness of sex-specific aging trajectory curves to the exclusion of individual age groups. Each plot shows the aging trajectories for male (teal) and female (reddish orange) samples in GTEx for a group of biological pathways (grouped according to KEGG hierarchy). The pathway targeting score on the y-axis is the mean in-degree of all genes in that group of pathways, aggregated across samples within each age group. The x-axis represents age. Each plot overlays 20 leave-one-bin-out LOESS curves (thin transparent lines), each fit after excluding all samples from one age group, alongside the main LOESS trajectory fit on the full dataset (thick line, ±95% confidence interval ribbon). The consistency of the leave-one-bin-out curves with the main trajectory across all pathway classes suggests that the observed transitions between ages 40 and 60 in females are not driven by any single age group or its sex composition.

**Figure S8:**
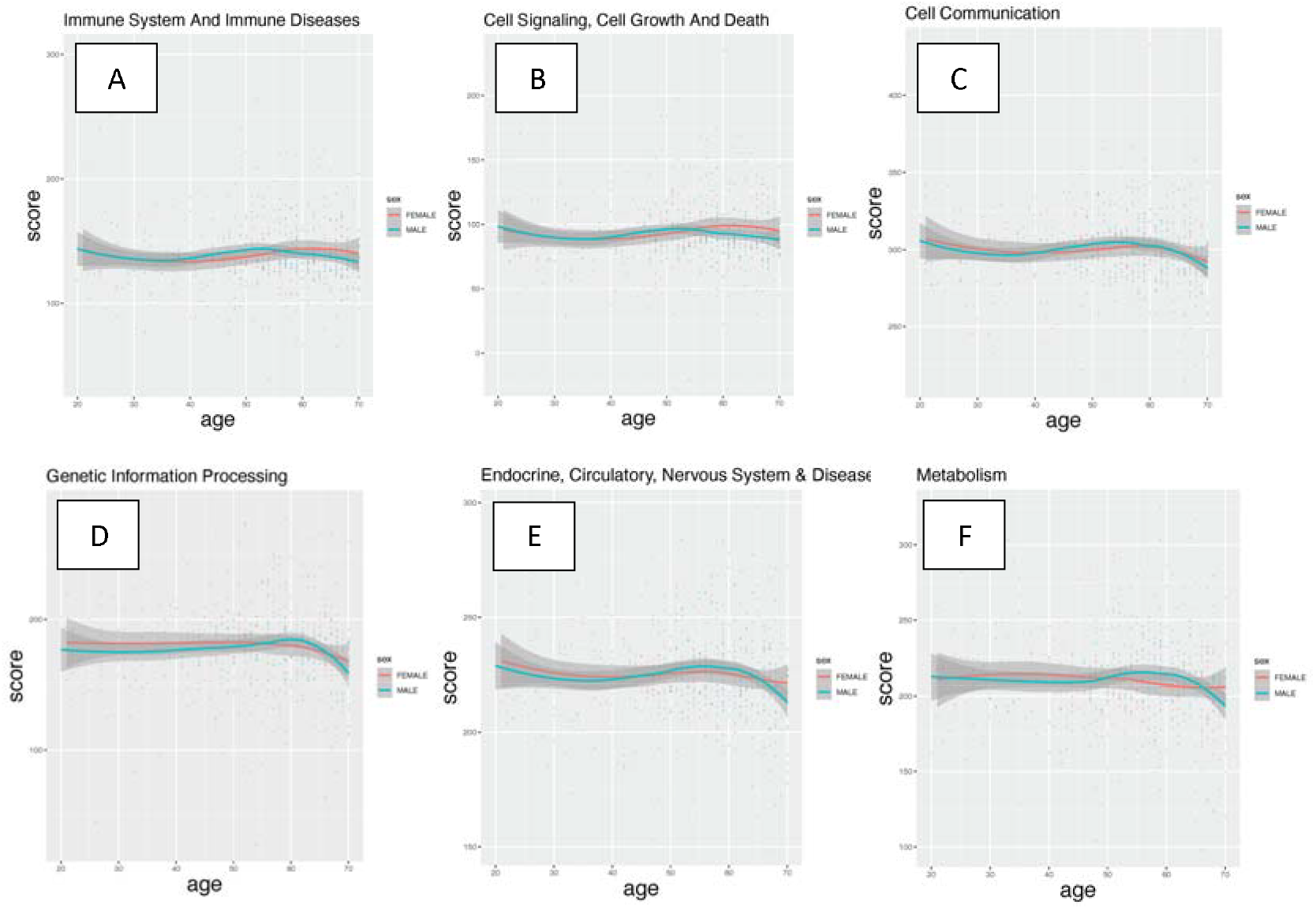
Effect of sex on aging pathway trajectory curves using raw age versus age-grouped values. Each plot shows the aging trajectories for male (teal) and female (reddish orange) samples in GTEx for a group of biological pathways (grouped according to KEGG hierarchy). The pathway targeting score on the y-axis is the mean in-degree of all genes in that group of pathways. When the x-axis represents raw age (top row), both sexes produce largely flat LOESS curves, as individual sampling fluctuations at each integer age dominate the signal and obscure underlying age-related trends. Aggregating samples into age bins before fitting (Figure 3) suppresses this noise and reveals the sex-specific aging trajectories, including the transitions between ages 40 and 60 observed in females.

**Figure S9:**
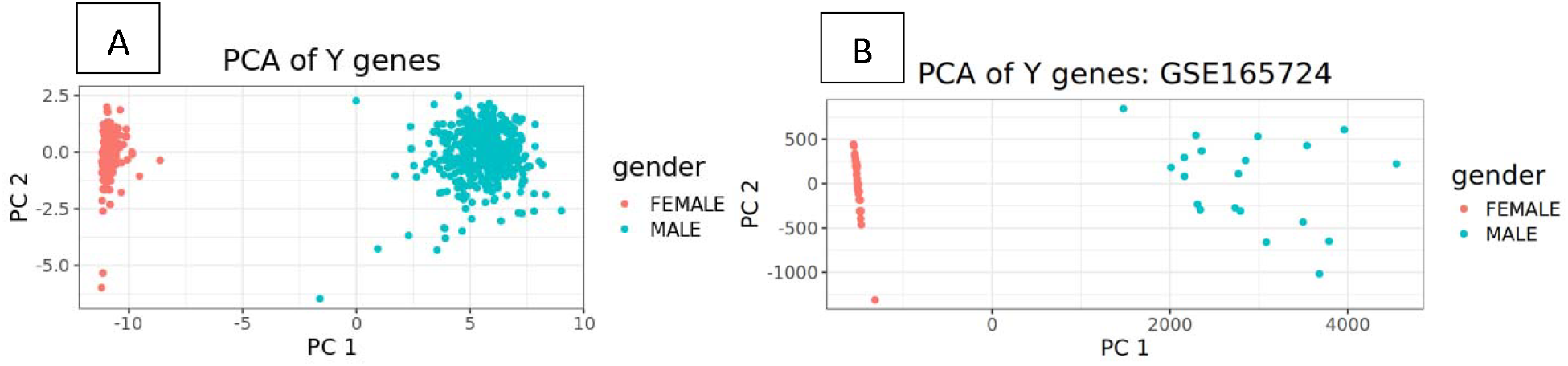
Reported gender aligns with chromosomal complement biological sex. In (A) GTEx and (B) GSE165724, we performed principal component analysis on the expression levels of the Y genes and drew scatterplots with the first two principal components (PC1 and PC2). Samples reported females and males in the original clinical data were marked orange and green, respectively.

**Figure S10:**
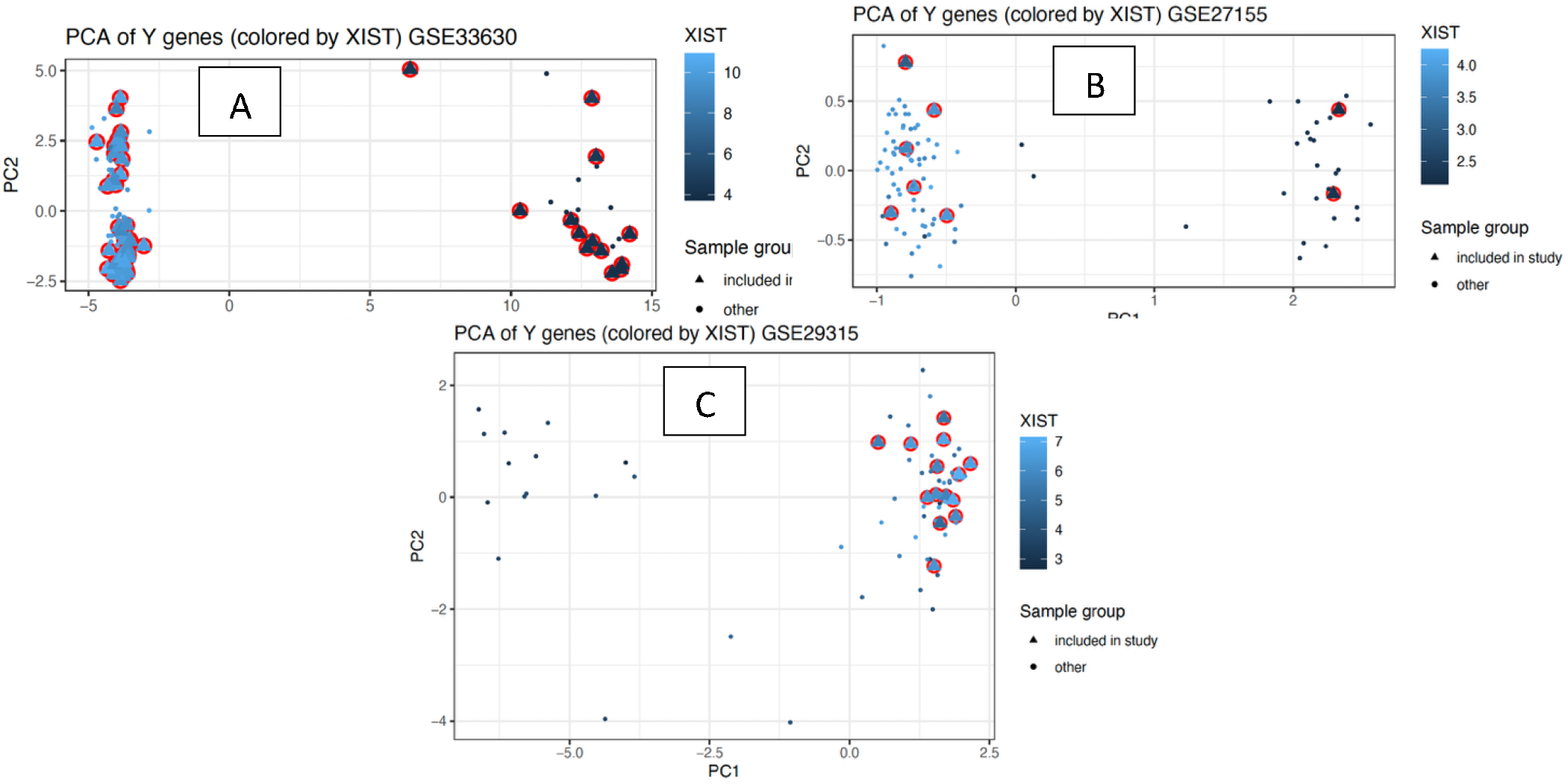
Biological sex identified by expression of genes on the Y chromosome and expression levels of XIST on the microarray datasets (A) GSE33630, (B) GSE27155 and (C) GSE29315. We performed principal component analysis on the expression levels of the Y genes and drew scatterplots with the first two principal components (PC1 and PC2). The datasets contained normal thyroid and samples from multiple diseases. We used all samples for sex identification but included only normal thyroid samples, Hashimoto’s Thyroiditis (HT) and Anaplastic Thyroid Carcinoma (ATC) in the study. Samples included in the study are denoted by triangles with a red round border. Other samples not included in the study are denoted by dots. The intensity of color represents expression levels of XIST gene, darker colored samples have lower expression (marked as males) and lighter colored samples have higher expression (marked as females).

